# Integrated RNA and metabolite profiling of urine liquid biopsies for prostate cancer biomarker discovery

**DOI:** 10.1101/599514

**Authors:** Bongyong Lee, Iqbal Mahmud, John Marchica, Paweł Dereziński, Feng Qi, Fubo Wang, Piyush Joshi, Felipe Valerio, Inoel Rivera, Vipul Patel, Christian P. Pavlovich, Timothy Garrett, Gary P. Schroth, Yinghao Sun, Ranjan J. Perera

**Affiliations:** Cancer and Blood Disorders Institute, Johns Hopkins All Children’s Hospital, 600 6th Avenue South, St. Petersburg, FL 33701 USA; Department of Oncology, Sydney Kimmel Cancer Center, Johns Hopkins University School of Medicine, 401 N. Broadway, Baltimore, MD 21287 USA; The James Buchanan Brady Urological Institute, Department of Urology, The Johns Hopkins University School of Medicine, 4940 Eastern Avenue, Baltimore, MD 21224, USA; Sanford Burnham Prebys Medical Discovery Institute, 6400 Sanger Road, Orlando, FL 32827 USA; Department Pathology, Immunology and Laboratory Medicine, University of Florida, College of Medicine, 1395 Center Drive, Gainesville, FL 32610 USA; Department of Inorganic and Analytical Chemistry, Poznan University of Medical Sciences, Grunwaldzka 6 Street, 60-780, Poznan, Poland; Department of Urology, Shanghai Changhai Hospital, Second Military Medical University, 168 Changhai Road, Shanghai, China; Florida Urology Associates, 1812 N. Mills Avenue, Orlando, FL 32803 USA; Global Robotics Institute, 410 Celebration Place, Suite 200, Celebration, FL 34747 USA; Illumina, Inc., 5200 Illumina Way, San Diego, CA 92122 USA

**Keywords:** Prostate cancer, metabolomics, biomarkers, gene expression, GOT1, glutamate

## Abstract

Sensitive and specific diagnostic and prognostic biomarkers for prostate cancer (PCa) are urgently needed. Urine samples are a non-invasive means to obtain abundant and readily accessible “liquid biopsies”. Herein we used urine liquid biopsies to identify and characterize a novel group of urine-enriched RNAs and metabolites in PCa patients and normal individuals with or without benign prostatic disease. Differentially expressed RNAs were identified in urine samples by deep sequencing and metabolites in urine were measured by mass spectrometry. The mRNA and metabolite profiles were distinct in patients with benign and malignant disease. Integrated analysis of urinary gene expression and metabolite signatures unveiled an aberrant glutamate metabolism and tricarboxylic acid (TCA) cycle node in prostate cancer-derived cells. Functional validation supports a role for glutamate metabolism and glutamate oxaloacetate transaminase 1 (GOT1)-dependent redox balance in prostate cancer, which can be exploited for novel biomarkers and therapies.

## Introduction

More than 180,000 men were diagnosed with prostate cancer (PCa) in the U.S. in 2016; 26,000 of these patients will die of the disease^1^. PCa is the second most frequently diagnosed cancer and the fifth leading cause of cancer deaths in men worldwide^2^. Although radiotherapy and surgery for localized PCa are effective, the prognosis for patients with advanced disease is poor. A test to detect PCa with high sensitivity and specificity at an early stage is a medical imperative. Moreover, there is an urgent need for novel therapeutic approaches to manage this insidious and prevalent disease.

Serum prostate specific antigen (PSA) levels have been used for PCa diagnosis and screening for over thirty years, and digital rectal examination (DRE) for even longer^3^. However, PSA has poor sensitivity and specificity and does not distinguish indolent from aggressive cancers^3,4^. Prostate cancer antigen 3 (PCA3), a prostate-specific non-coding RNA, was approved by the FDA in 2012 as the first PCa molecular diagnostic test for a specific clinical indication (need for repeat prostate biopsies in men aged >50 years with suspected PCa based on PSA levels and/or DRE and/or one or more previous negative biopsies)^5^. However, the value of the PCA3 test is limited by significant individual variability, better performance in the repeat biopsy setting, and conflicting data on the relationship between score and cancer grade using the most common threshold of 35^6^. Hence, there is a dire need for novel molecular diagnostic tools to more accurately detect and predict the behavior of localized PCa.

The kidneys produce urine to eliminate soluble waste from the bloodstream. Urine is an abundant biofluid for molecular or cellular analyses and is useful in the diagnosis and management of bladder, ovarian, and kidney diseases^7–9^. Urine contains over 2500 metabolites^10^ and provides a window through which to view cellular biochemical reactions and intermediary metabolism. The metabolite signature in urine will reflect the impact of gene regulation, enzyme activities, and alterations in metabolic reactions occurring in the different cell types found along the urogenital tract.

Cancer cells exhibit perturbed metabolism that enables proliferation and survival^11^. Therefore, metabolomic profiling has been a fruitful approach for the identification of early cancer biomarkers^12,13^. Furthermore, certain metabolic states are associated with prognosis in advanced cancers^13^. Several metabolomics studies have revealed PCa-specific metabolic phenotypes in serum, tissue, and urine^14,15^. Indeed, an intermediate metabolite of glycine synthesis and degradation, sarcosine, has been described as a putative PCa biomarker in urine^16^. However, the utility of sarcosine as a biomarker is controversial and clinical validation has been elusive^17^.

Metabolomics data have been integrated with comprehensive gene expression analyses to better interrogate complex gene and metabolic networks. Integrating multiple aspects of biological complexity using different unsupervised approaches can help to pinpoint the most important and reproducible pathways driving biological processes and hence reveal robust biomarkers or promising drug targets^18,19^. Here we performed metabolite profiling and high-throughput RNA sequencing of urine from patients with benign prostatic hyperplasia, prostatitis, and PCa. Our aims were to (a) discover cancer-specific changes in the urine with utility as sensitive and specific PCa biomarkers either alone or in combination and (b) identify novel drug targets for PCa. Importantly, our approach used single void urine samples (i.e., without prostatic massage) as proof-of-principle of how a simple urine specimen can be used for biomarker and target discovery. Integrated analysis of metabolomic and transcriptomic data from these liquid biopsies revealed a glutamate metabolism and tricarboxylic acid cycle node that was specific to prostate-derived cancer cells and cancer-specific metabolic changes in urine. Functional validation *in vitro* provided mechanistic support for a pivotal role for GOT1-dependent glutamate metabolism in redox balance and cancer progression.

## Results

### Deep sequencing of urine-secreted mRNAs

Normal voided urine from men contains small numbers of exfoliated cells from different parts of the urinary tract including urothelial cells, squamous cells, renal tubular cells, and glandular cells including prostate epithelial cells^20^. PCa cells are shed into urine and can be successfully isolated, processed, and analyzed by various molecular techniques^3^, thereby providing a rich substrate for biomarker detection. We sought to exploit this readily accessible and copious substrate from PCa patients for biomarker discovery and, in turn, elucidate novel mechanistic aspects of PCa.

Quality output from current next-generation sequencing (NGS) technology depends on the availability of high-quality RNA. An initial challenge was that the quality and quantity of RNA extracted from the very small number of exfoliated cells in urine was poor^21^ (**Supplementary Figure 1**). To overcome the problem, we performed sequence-specific capture (Illumina TruSeq RNA Access) with the urine samples to reduce ribosomal RNA and enrich for exonic RNA sequences. With this approach, we successfully sequenced 11 PCa (for clinical details, see **Supplementary Table 1**), 12 normal, and one pooled set of three normal samples (combined due to individually low RNA yields). The 3825 RNA transcripts that were detected in 20 samples readily but not perfectly segregated into normal and PCa groups (**Supplementary Figure 2**). We concluded that RNA expression analysis of urine liquid biopsies by itself was unlikely to reveal sensitive and specific PCa biomarkers.

We next identified cancer-specific gene signatures. Among 5510 differentially expressed transcripts, 4662 had reads per kilobase of transcript per million mapped reads (RPKM) values greater than one, and 116 transcripts (110 genes) were significantly up- or downregulated in PCa (Table 1). Known PCa markers were upregulated in PCa urine (**Supplementary Table 2**), and differentially expressed genes were enriched for a number of important cancer pathways including PCa signaling, molecular mechanisms of cancer, PI3K/AKT signaling, and NF-κB signaling (**Supplementary Table 3**). To our knowledge, this is the first time that RNA-seq has been successfully applied to urine samples to profile coding genes.

**Table 1.**
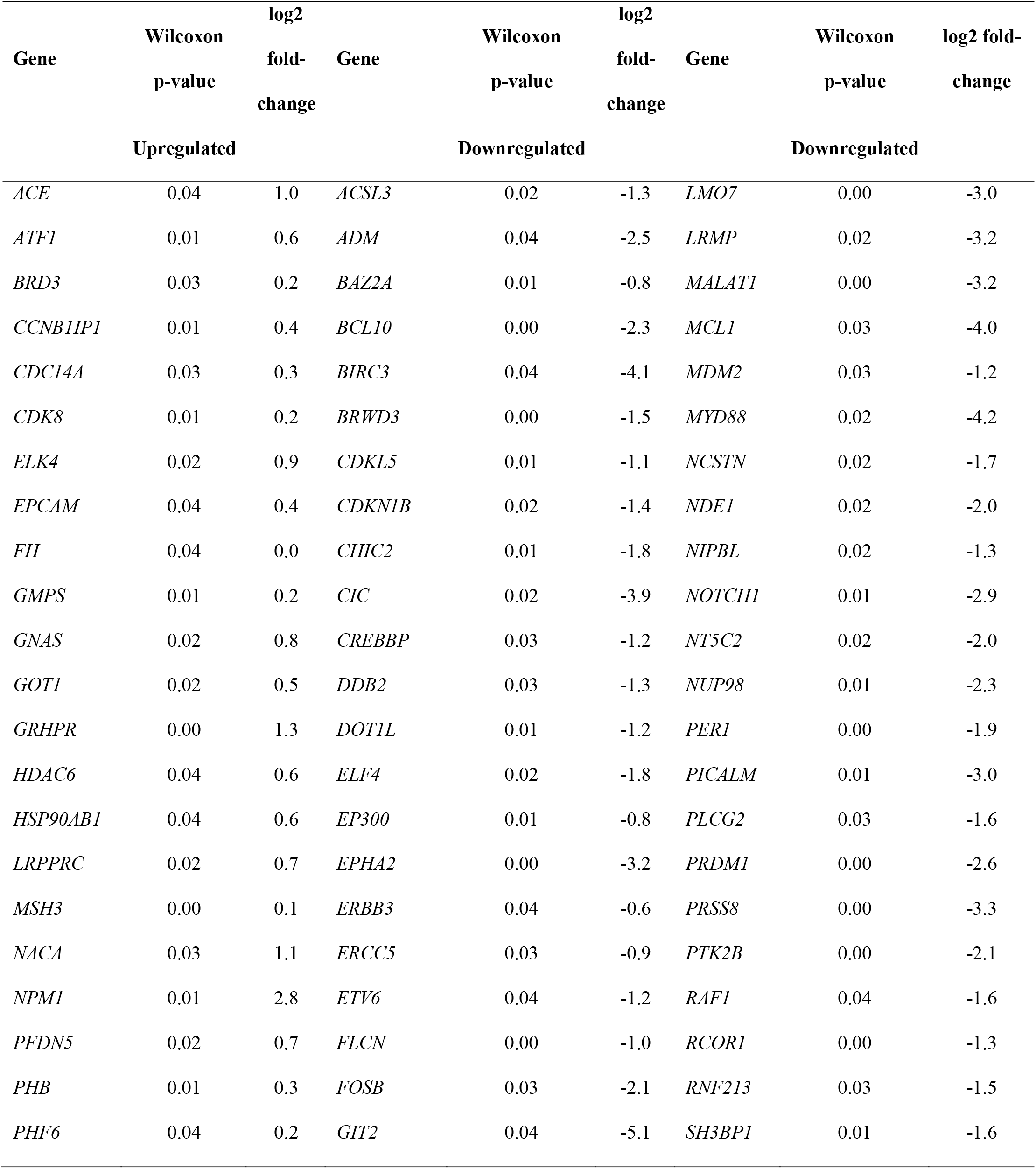
Differentially expressed genes in prostate cancer urine samples.

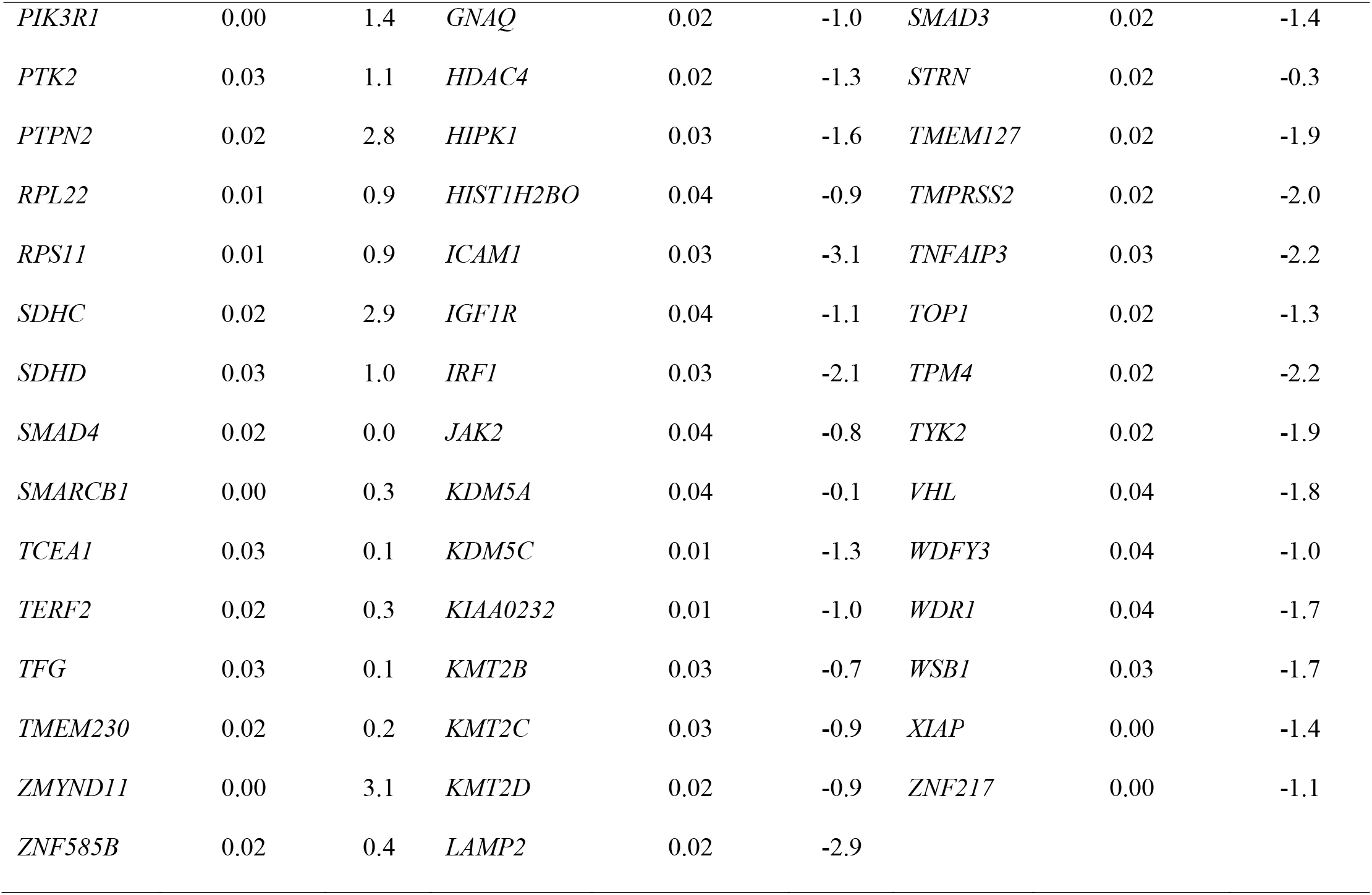

Thirty-seven genes were significantly upregulated in PCa urine samples (Table 1 and Figure 1a) compared to normal urine samples. To bolster confidence that the RNA originated from the patient’s cancer rather than contaminating cells, we examined their expression in The Cancer Genome Atlas (TCGA) data (Figure 1b and **Supplementary Table 4**). Of these 37 genes, 35% (13/37) were significantly upregulated in primary tumors compared to normal (Figure 1b **and** 1c). Three of these genes were transcription factors (*ELK4*, *SMARCB1*, *BRD3*) and six were known oncogenes (*TFG*, *NACA*, *BRD3*, *ELK4*, *NPM1*, *RPL22*)^22^. When quantified in two representative PCa cell lines (LNCaP and PC3), most transcripts were upregulated in both cell lines compared to normal prostate epithelial cells (PrEC) except for *NACA* (downregulated in both cell lines), *BRD3* and *EPCAM* (decreased in PC3 cells), and *HDAC6* (downregulated in LNCaP cells) (Figure 1d). Gene set enrichment analysis (GSEA) revealed thirteen overrepresented pathways in PCa urine compared to normal urine samples based on normalized gene set enrichment scores (NES) (Figure 1e). Among them, the TCA cycle and alanine, aspartate, and glutamate metabolism were significantly enriched (Figures 1f **and** 1g). This finding is highly consistent with current knowledge of prostate cancer disease progression using tissues and other biofluids^16,22^. Taken together, our data suggest that the transcriptional profiles generated from cells residing in urine from PCa patients are likely to originate from cancerous prostate epithelial cells rather than other urinary tract contaminants.

**Figure 1.**
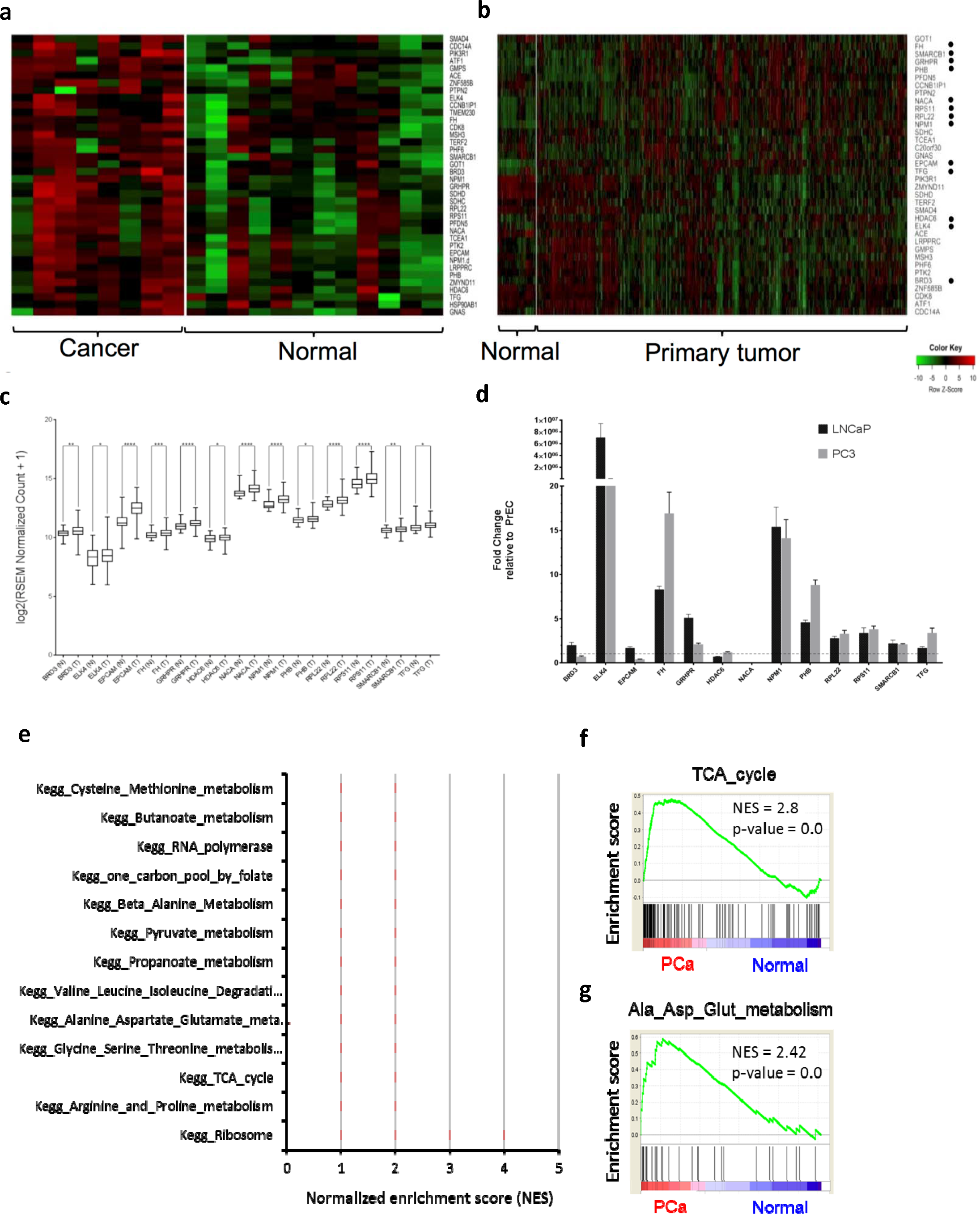
RNA-seq of cells extracted from the urine of patients with (n=8) and without (n=12) prostate cancer and validation of their differential expression in The Cancer Genome Atlas (TCGA) data. **(a)** A heatmap showing expression of 37 significantly upregulated genes in cancer liquid biopsies. **(b,c)** Among 37 upregulated genes, 13 genes (*FH, SMARCB1, GRHPR, PHB, NACA, RPS11, RPL22, NPM1, EPCAM, TFG, HDAC6, ELK4, BRD3*) were significantly upregulated in primary tumors (n=497) compared to normal (n=52) in TCGA data. In the heatmap, black dots next to the gene name mark the genes upregulated in primary tumor compare to normal in TCGA data. The TCGA project for PCa data is publicly available for download at https://portal.gdc.cancer.gov/projects/TCGA-PRAD. **(d)** These 13 genes were also tested in two prostate cancer cell lines (LNCaP and PC3), and most were overexpressed apart from *NACA*, which was downregulated in both cell lines; *BRD3* and *EPCAM*, which were decreased in PC3 cells, and *HDAC6*, which was downregulated in LNCaP cells. **(e)** Normalized Enrichment Score plot of the top 13 pathways in cancer urine. **(f)** Geneset Enrichment Score Plot (GSEA) of the TCA cycle (NES = 2.8, p-value = 0.0). **(g)** GSEA of the alanine, aspartate, and glutamate metabolism (NES = 2.42, p-value = 0.0). GSEA was conducted using GSEA software from the Broad Institute (http://software.broadinstitute.org/gsea/index.jsp).

### Validation of urine gene signatures in tumor tissue

Since exfoliated cells in urine represent a mixture of cell types, we next established that the gene expression profiles of urine-exfoliated cells represented expression in prostate tissue using RNA-seq data for 65 PCa and matched normal prostate tissues^23^ (Table 1). Most of the 110 up- or downregulated genes in urine exfoliated cells agreed with tissue gene expression (Figure 2 **and Supplementary Table 5**). Of 37 upregulated genes, 34 genes were upregulated in PCa tissue, and 27 out of 34 genes (79%) were significantly upregulated (Figure 2). For the downregulated genes, 46 out of 73 genes (60%) were downregulated in PCa tissue compared to normal tissue (**Supplementary Table 5**). These results suggest that the PCa gene signature detected in urine exfoliated cells represents a *bone fide* PCa signature, especially with regard to upregulated genes, which may therefore represent more robust biomarkers.

**Figure 2.**
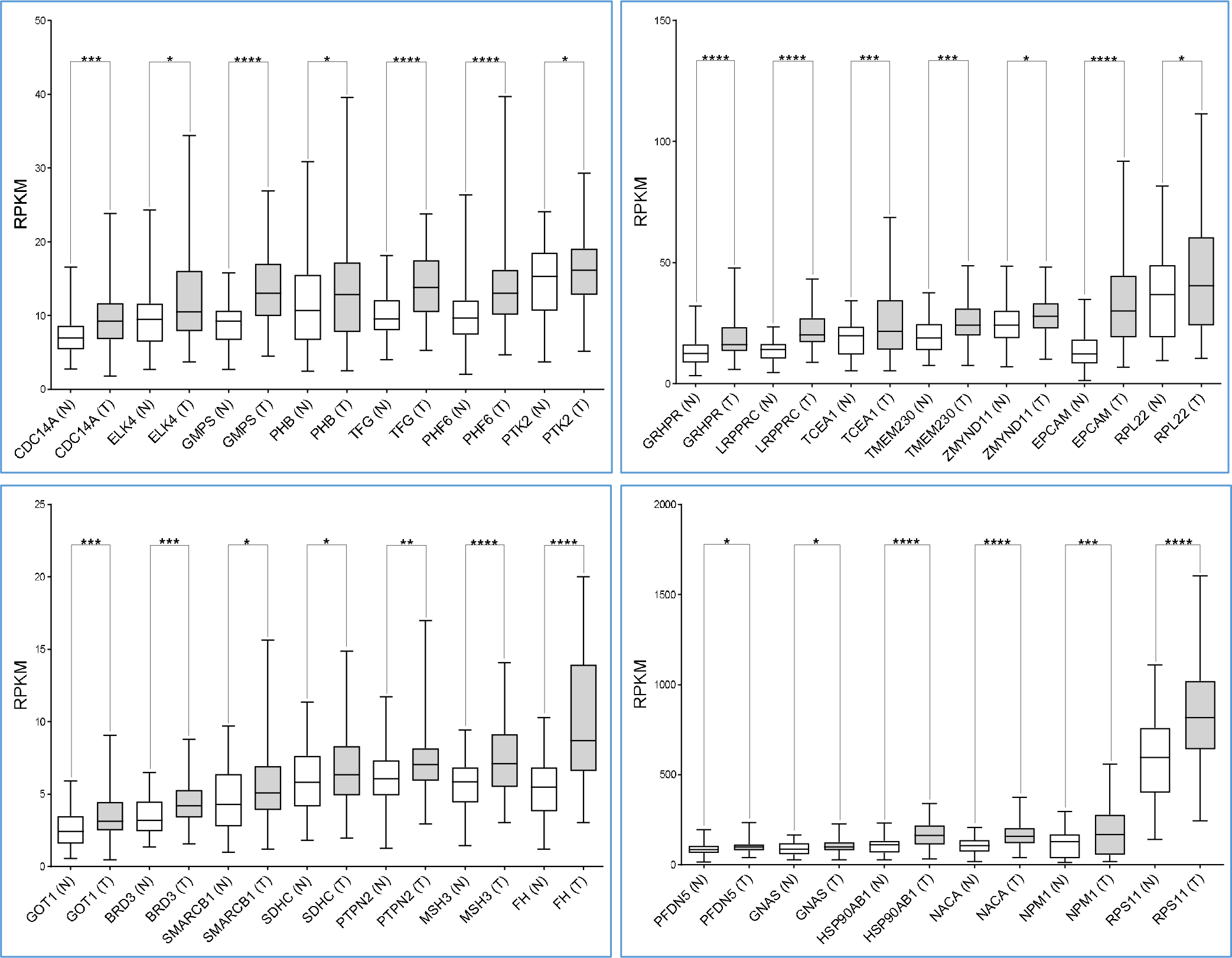
Validation of the expression of 37 genes in prostate tumor tissues. The 37 gene signature from PCa patient urine RNA was confirmed in 65 pairs of tumor and adjacent normal tissue RNA-seq data. Thirty four out of 37 genes were upregulated in PCa tumor tissue. Among them, 27 genes were significantly upregulated (Student *t*-test, * p<0.05, ** p<0.01, *** p<0.001, **** p<0.0001).

Thus, we performed principal component analysis (PCA) of the 37 upregulated genes in the 65 patient tissue RNA-seq data. This 37-gene signature divided the tumor samples into two distinct groups, A and B (Figure 3a), which did not differ with respect to Gleason score, tumor stage, or metastasis status (**Supplementary Figure 3**). However, the two groups did show significant differences in *PCA3* and *KLK3* expression (**Supplementary Figure 4**), being significantly higher in group B than in group A patients (**Supplementary Figure 4**).

**Figure 3.**
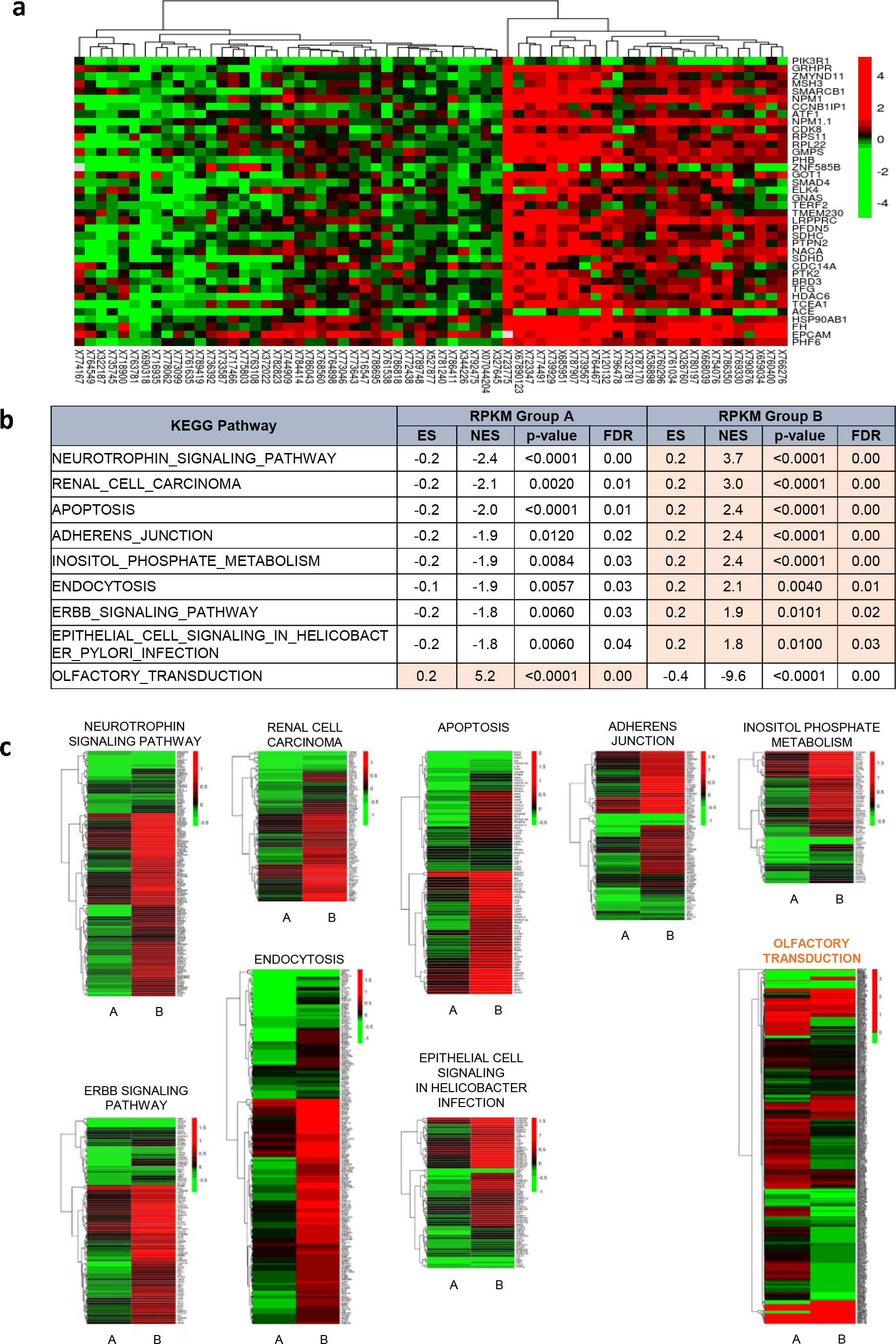
The urine 37 gene signature distinguishes PCa patients into two groups. **(a)** Unsupervised clustering of log-fold change (logFC) values shows that patients can be grouped into two groups, Group A and Group B, exhibiting lower and higher expression of the 37 genes compared to the matched normal, respectively. The logFC was obtained from the log2 ratio of RPKM values of tumor and matched normal of the listed genes. **(b,c)** KEGG pathway analysis suggests that the two molecularly identified groups are physiologically distinct, as shown by opposite enrichment of the depicted pathways. The logFC values were obtained from comparing tumor versus matched normal samples from each group via paired limma analysis.

With respect to pathway differences between groups, nine pathways were significantly different: eight pathways were significantly upregulated (FDR>0.05) in group B, and one pathway, olfactory transduction, was upregulated in group A (Figure 3b **and** 3c). Therefore, the 37-gene signature in urine samples represents prostate tissue gene expression and might be useful to distinguish advanced PCa (higher *PCA3* and *KLK3* levels in cancer) as well as to detect PCa itself.

### Metabolomic profiling of urine from normal subjects and patients with diseased prostates

Targeted or global strategies have been used to profile metabolites in urine samples and identify PCa biomarkers^14,16,24,25^, but results have been highly variable^26^. In the first unbiased metabolomics study measuring 1126 metabolites in 262 clinical samples including 110 urine samples, the glycine derivative sarcosine was elevated in PCa tissue and urine from PCa patients, and functional validation of the oncogenic role of sarcosine was provided *in vitro*^16^. However, sarcosine was not a reproducible prognostic marker in independent cohorts^15,17^, a common finding in single-biomarker studies that possess neither the specificity nor sensitivity for clinical development^27^.

With this in mind, we performed global metabolite profiling of urine from patients with normal prostates, benign prostatic hyperplasia (BPH), prostatitis (PTT), and PCa to discover cancer-specific metabolic changes. In global metabolite profiling, the metabolic profiles of urine specimens from normal subjects and patients with cancer were distinct and separate by PCA (Figure 4a **and** 4b), whilst there was significant overlap between the profiles obtained from patients with BPH and PTT (Figure 4c). Positive and negative ion data were first normalized to the specific gravity and then normalized to the total ion signal for all subsequent statistical analyses (**Supplementary Figure 5a**). Positive and negative ion data sets were treated separately, and initial analysis was performed with PCA. The negative ion data set separation by PCA was very distinct between PCa and control groups, with BPH and PTT clustering together but as a separate cluster from PCa and control. Separation was primarily observed along PC1 (Figure 4a). However, no correlation was observed between PSA scores and metabolic profiling. This result was reproducible with a second set of urine samples (**Supplementary Figure 5b**). Further, subsequent hierarchical clustering-based heatmap analysis revealed distinctly higher abundance of global metabolites in PCa urine samples compare to normal samples (**Supplementary Figure 5c**); over 180 metabolites in positive mode and 140 metabolites in negative ion mode were detected from the extracted urine samples, respectively.

**Figure 4.**
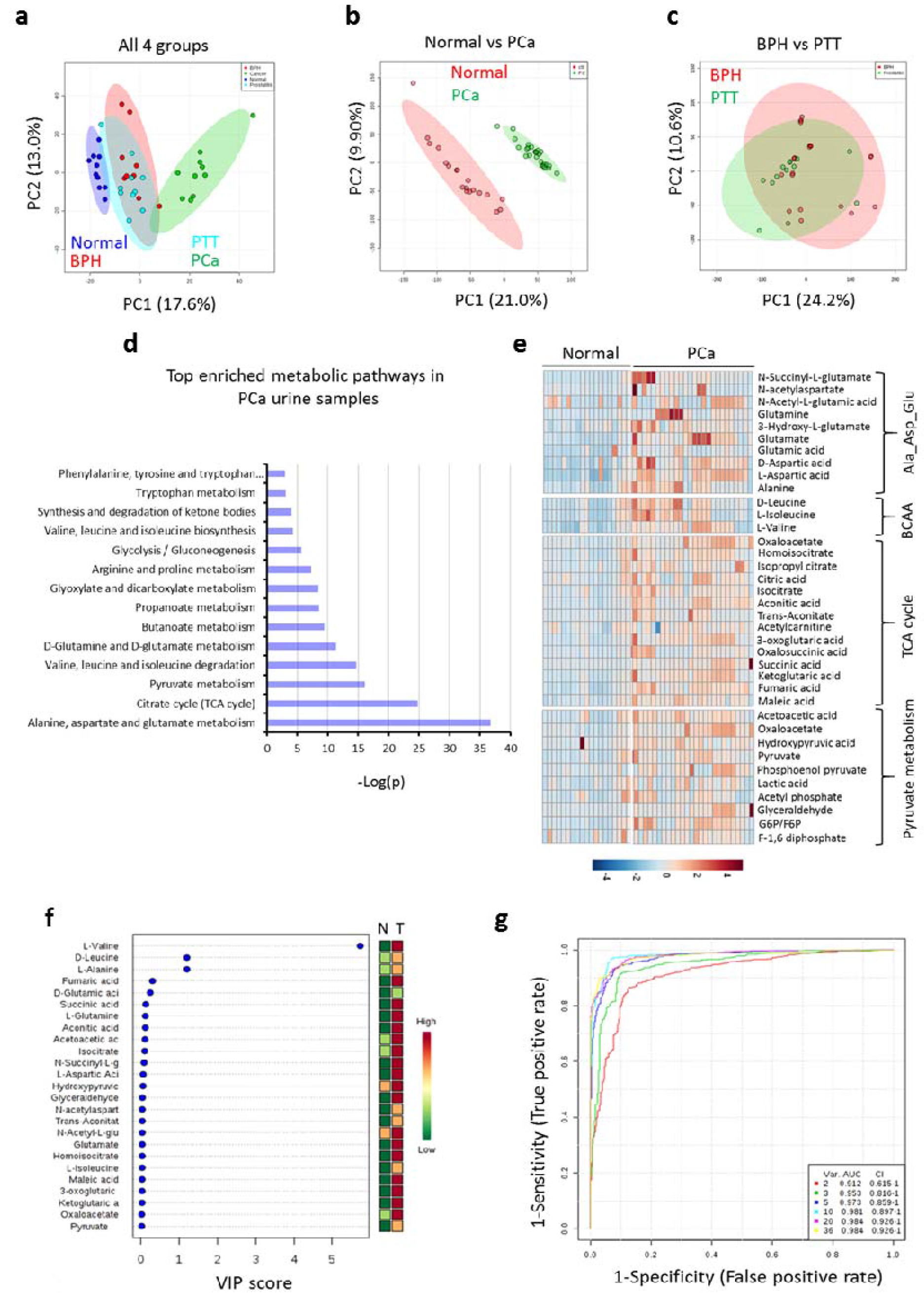
Global untargeted urine metabolomics profile data reveals distinct metabolic differences between PCa and normal. **(a)** Multivariate principal component analysis (PCA) scores plot for normal (n=20), BPH (n=20), PTT (n=11), and PCa (n=20). **(b)** PCA score plot for BPH and PTT. **(c)** PCA score plot for normal and PCa. **(d)** Heat map of the differential metabolites in PCa tissue. Color bar indicated the relative abundance of the metabolites, with red indicating a higher concentration and blue indicating a lower concentration. Ala_asp_Glu, Alanine, Aspartate, and Glutamate metabolism; BCAA, Branched Chain Amino Acid metabolism, G6P, Glucose-6-phosphate; F6P, Fructose-6-phosphate. P < 0.05 was considered significant. **(e)** List of significant metabolic pathways in PCa urine samples. The p-value is shown in negative log10 scale. **(f)** Metabolites ranked by their contributions and shown as variable importance in the projection (VIP) scores. **(g)** ROC curves for the predictive model. Shown as combination metabolite models calculated from the logistic regression analysis.

Comprehensive metabolic networks in PCa urine samples have not been well studied, and understanding differential metabolic pathway utilization in PCa might contribute towards the development of robust biomarkers. Therefore, we performed metabolite-based pathway enrichment analysis (**Supplementary Figure 5d**), which revealed 14 significantly impacted metabolic pathways in the urine metabolome (Figure 4d). Notably, glutamate metabolism, TCA cycle metabolism, pyruvate metabolism, and several amino acids pathway metabolites were identified at higher levels in PCa urine samples compare to normal (Figure 4e), consistent with previous tissue-based studies^16,23^.

We next conducted metabolite-based urinary biomarker screening, with metabolite contribution assessed by examining the variable importance in projection (VIP) score, which is calculated from the weighted sum of the square for each partial least square design (PLS) loadings for each principal component. Of the top twenty five variables identified by VIP scores, all were metabolite variables that significantly contributed to the class separation of normal and PCa samples (Figure 4f). We then conducted multivariate receiver operating characteristics (ROC) curve-based exploratory biomarker analysis to identify a diagnostic PCa-related metabolite signature. To better predict PCa, the top 50 discriminatory metabolites were identified via logistic regression (Figure 4g). A combination of six metabolites showed better discrimination (AUC>98%) than each metabolite individually (AUC<91%) (Figure 4g): aconitic acid (AUC=0.97), succinic acid (0.96), fumaric acid (AUC 0.955), oxaloacetate (AUC=0.952), α-ketoglutaric acid (AUC=0.921), and glutamate (AUC=0.951) (**Supplementary Figure 6a**).

Given that Gleason score (GS) status correlated with PCa tumor progression, an additional independent 11 normal, 11 GS-6, 11 GS-7, 11 GS-8, and 11 GS-9 PCa urine samples were collected and again subjected to global metabolomics analysis using mass spectrometry (Figure 5). Metabolite data were analyzed using same statistical approaches as in Figure 4. Furthermore, metabolome data accuracy were validated by PLS-DA-based Q2 model (Figure 5a). Supervised multivariate statistical analysis of the global metabolome revealed a profound trend of clustering with respect to the four different GS groups and the normal urine samples (Figure 5b). Hierarchical clustering heatmap analysis identified distinct metabolic signatures among normal and different GS groups (Figure 5c). Differential metabolite analysis revealed that most metabolites that were discriminatory across the normal and different GS group samples were involved in TCA cycle and glutamate metabolism (Figure 5d-i). Notably, level of these metabolites were significantly increased with highest GS, the aggressive form of PCa (Figure 5d-i).

**Figure 5.**
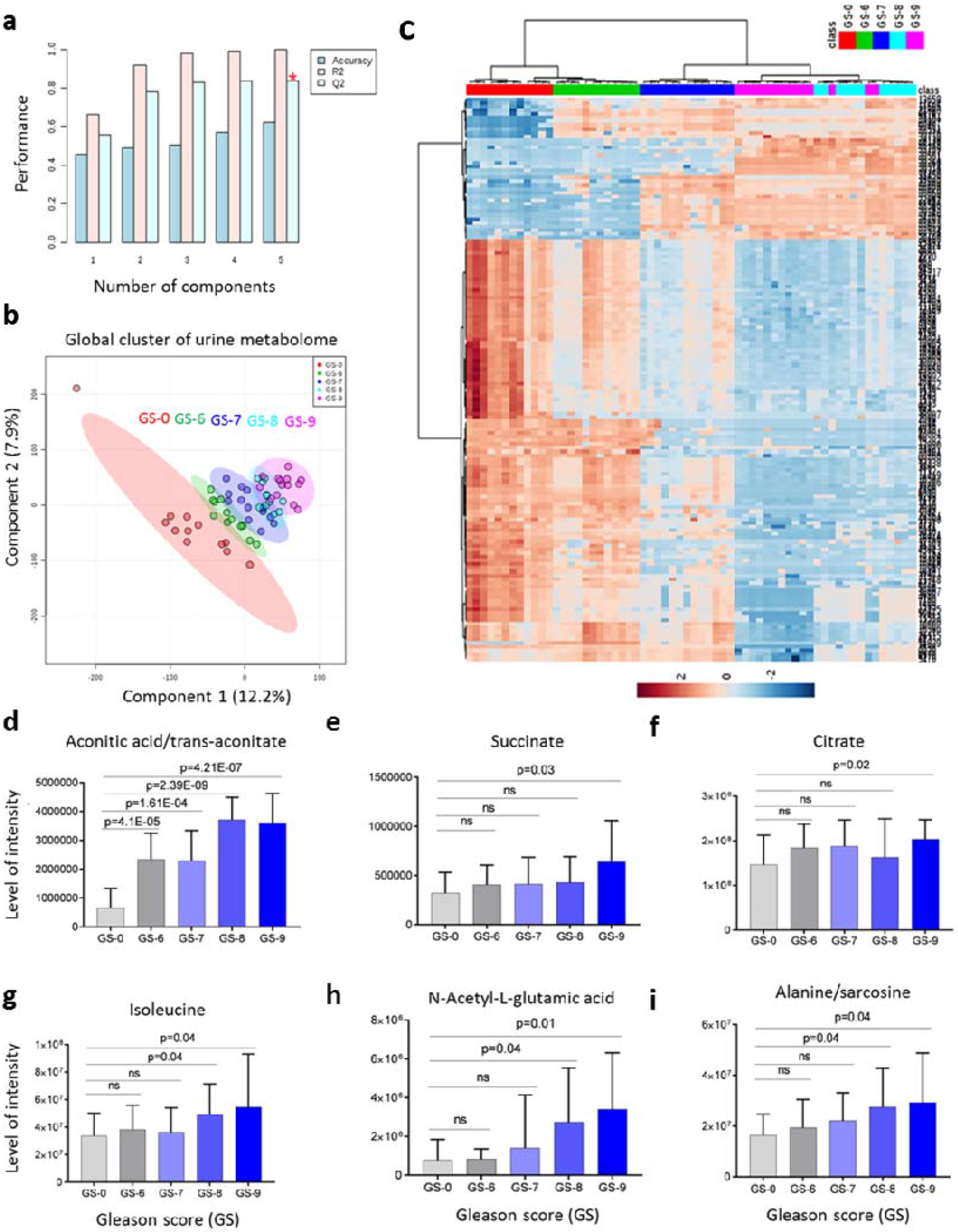
Correlation of TCA cycle and glutamate metabolism pathway metabolite levels with PCa GS (Gleason score). **(a)** Validation of PLS-DA (partial least square-differential analysis) model. Q2 indicates the accuracy of the model is over 80% when it includes top five components. (b) Plot of PLS-DA component 1 and component 2 for log2 autoscaled metabolite abundance data for normal versus different PCa GS. Circle indicate the 95% confidence interval of each sample group. (c) Heatmap showing the autoscaled abundances of global metabolome in normal versus different PCa GS-based urine samples. Data were analyzed using Euclidean distance calculation and ward clustering algorithm. (d-i) Correlation of representative TCA cycle and glutamate metabolism pathway metabolites levels with PCa Gleason score (GS). Statistical significance was analyzed by Student‘s t-test with two-tailed unequal variance.

Prostate carcinogenesis is known to involve metabolic reprogramming to provide sufficient energy for rapid cellular proliferation^28,29^. Many cancer cells exhibit augmented aerobic glycolysis, known as the Warburg effect, even in high-oxygen environments^30^. This metabolic adaptation helps provide essential cellular components such as lipids and nucleotides to support the anabolic needs of rapidly proliferating tumor cells. Beyond the Warburg effect, the TCA cycle and oxidative phosphorylation also play important roles in PCa^29,31^. Prostate epithelial cells normally produce certain components of prostatic fluid such as citrate, PSA, and polyamines^31^. Increased citrate production by prostate cells means that they favor citrate synthesis over citrate utilization. However, PCa cells degrade citrate and accumulate oxidized citrate, resulting in more efficient energy production^29,32^.

### Integrated gene expression and metabolite analysis

We reasoned that integrating changes in gene expression and metabolite levels evident in the urine samples would better reveal the key pathways driving PCa and hence pinpoint the most robust biomarkers. The integrated pathway analysis module of MetaboAnalyst^33^ was used to map both genes and metabolites to KEGG pathways to determine not just overrepresented pathways, but also the relative importance of the genes and compounds based on their relative locations (topology). The top three pathways most significantly enriched for differentially expressed genes and metabolites were: aminoacyl-tRNA biosynthesis; Ala, Asp, and Glu metabolism; and the TCA cycle) (p<0.001; Figure 6a and **Supplementary Table 6**). Aminoacyl-tRNA biosynthesis probably represents an increase in global protein translation and demand for protein synthesis in cancer cells^34^. However, Ala, Asp, and Glu metabolism and the TCA cycle are closely related pathways that are critical for energy generation and carbon and nitrogen metabolism for biomass accumulation^28^, especially in rapidly dividing cells such as cancer cells.

**Figure 6.**
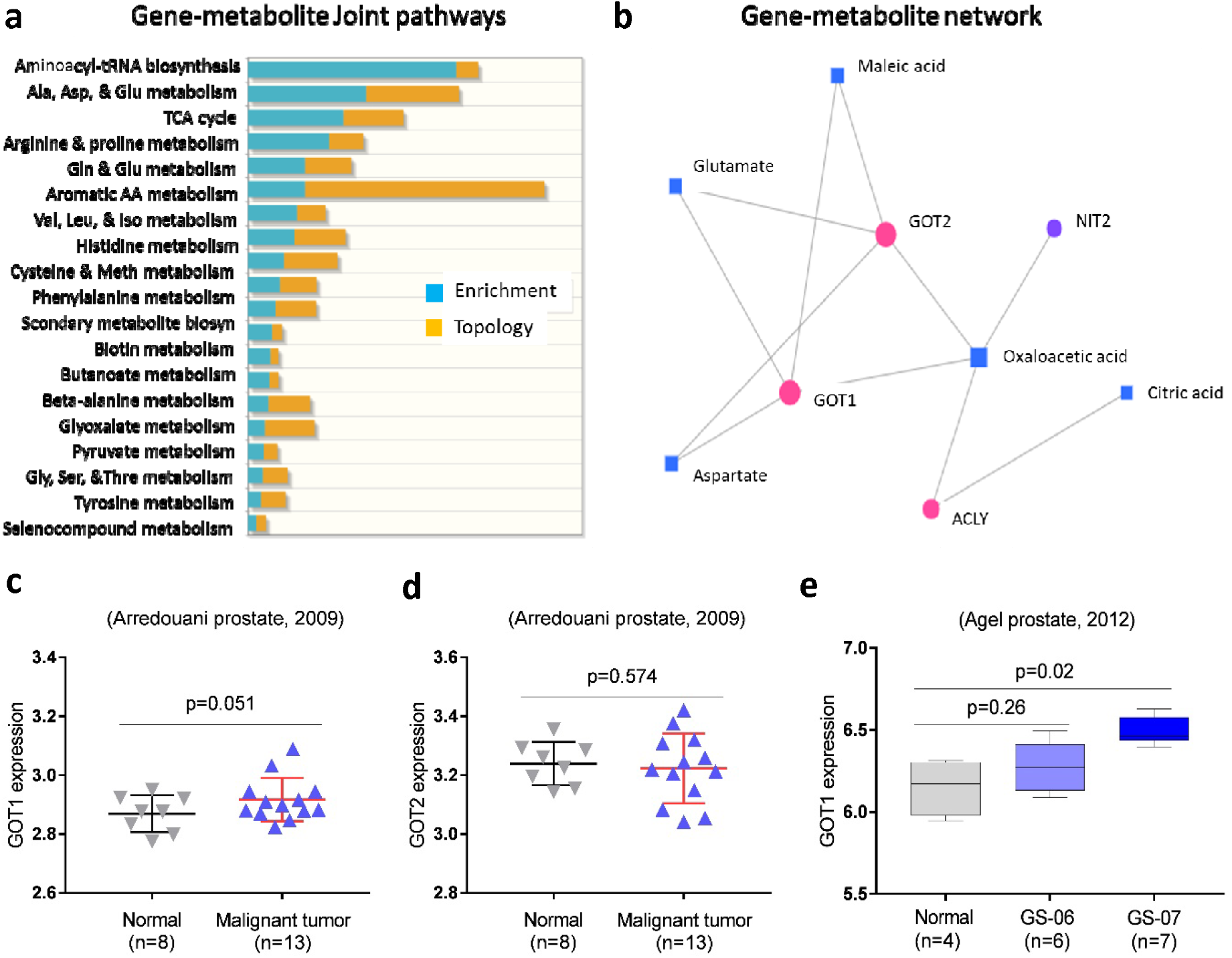
Integrated metabolic pathway enrichment analysis. **(a)** Gene-metabolite joint pathways were identified using MetaboAnalyst integrated pathway analysis module. **(b)** Gene-metabolite network exploration identified a GOT1- and GOT2-mediated interactive network that might influence the TCA cycle and other metabolic pathways. **(c, d)** Box plots of GOT1 and GOT2 expression in malignant prostate tumor compare to normal (data sources GSE55945). **(e)** A box plot of GOT1 expression across the prostate cancer Gleason score (GS) (data sources GSE30521).

Analysis of the top 25 metabolites identified by UHPLC-HRMS and their corresponding genes from RNA-seq revealed that GOT1- and GOT2-mediated metabolism was the main gene-metabolite interactive node influencing Ala, Asp, and Glu metabolism and the TCA cycle metabolism (Figure 6b **and Supplementary Figure 6**). In several PCa clinical datasets, GOT1 expression was significantly higher in malignant prostate compared to normal, whereas GOT2 expression showed no significant differences (Figure 6c **and** 6d). Interestingly, GOT1 expression was significantly elevated in high Gleason score tumors compared to controls (Figure 6e). Therefore, GOT1-mediated glutamate metabolism might be critical for PCa disease progression, and a better understanding of GOT1-driven metabolism could reveal a potential drug target and biomarker for PCa.

### Glutamate metabolism contributes to the cancerous phenotype via *GOT1*-mediated redox balance

*GOT1*, a cytosolic transaminase that converts Asp to Glu, and other genes involved in Gln metabolism such as *GLUD1*, *GOT1*, *GOT2*, and *MDH1* were significantly upregulated in PCa urine samples (**Supplementary Figure 7** and **Supplementary Table 7**). To investigate GOT1’s role as a regulatory metabolic node in prostate cancer, we knocked down *GOT1* in the prostate cancer cell lines LNCaP and PC3 using siRNA (Figure 7a). As expected, *GOT1* knockdown upregulated the upstream metabolites (**Supplementary Figure 7 and 8**) Glu [1.2-fold (LNCaP; *p*=*0.01*) and 1.4-fold (PC3; *p*=*0.03*)] and Asp (1.5-fold (LNCaP; *p*=*0.0004*) and 2.6-fold (PC3; *p*=*0.0006*)] in both cells lines. *GOT1* knockdown significantly decreased the viability of both LNCaP and PC3 cells (Figure 7b), consistent with previous reports that GOT1 repression suppresses tumor growth^35,36^ and the invasiveness and colony forming ability of PC3 cells (Figure 7c **and** 7d). We therefore examined the mechanism by which *GOT1* regulated prostate cancer cell viability.

**Figure 7.**
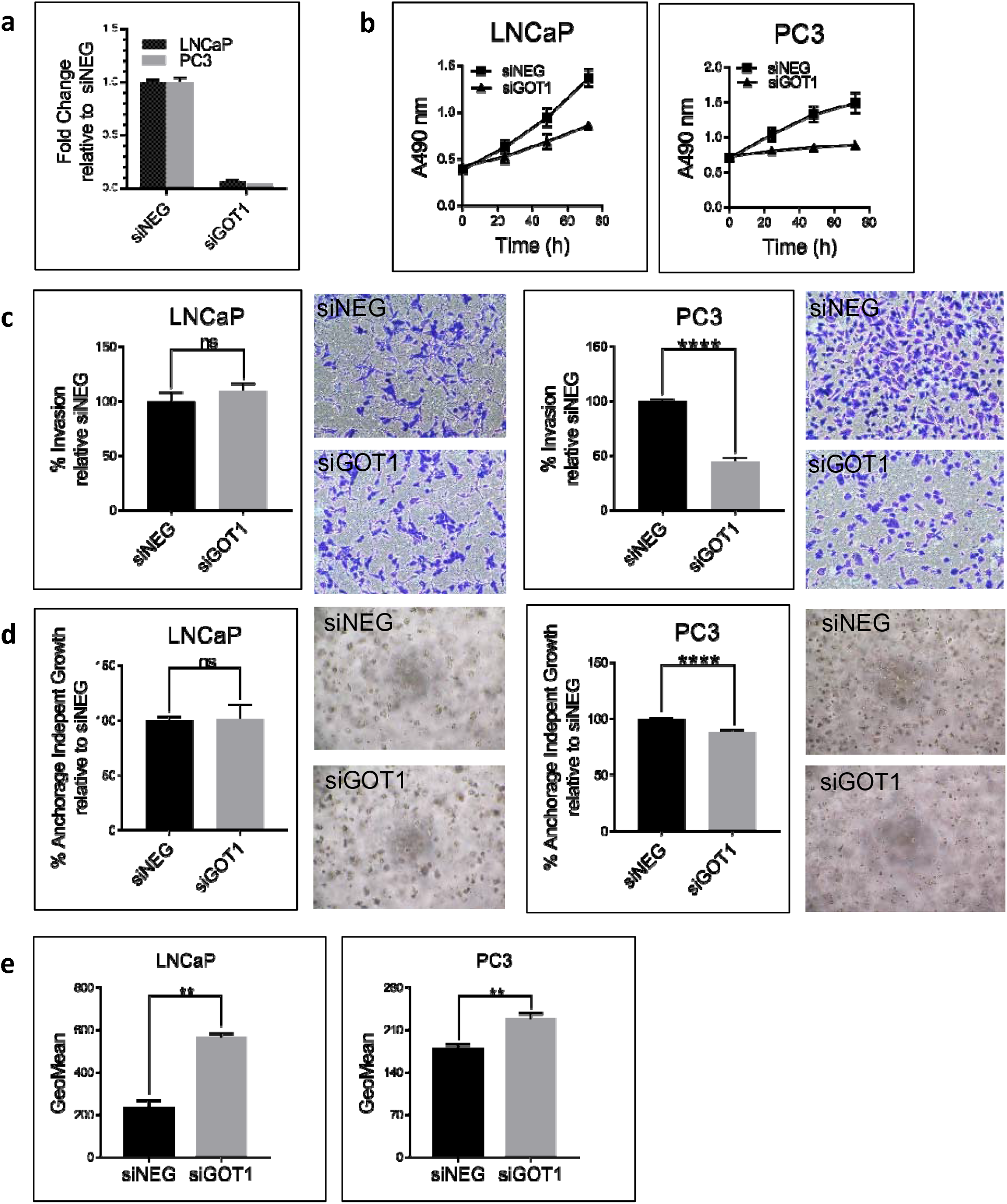
GOT1 supports proliferation, invasion, and colony formation in prostate cancer cell lines. **(a)** GOT1 knockdown in LNCaP and PC3 prostate cancer cell lines. **(b)** GOT1 knockdown significantly inhibits cell viability in PC3 and LNCaP cells. **(c)** Invasion and **(d)** anchorage-independent growth in prostate cancer cell lines upon GOT1 knockdown. **(e)** GOT1 knockdown significantly increases ROS production in PC3 and LNCaP cells. The data from three independent experiments were expressed as mean ± SD.

Maintaining NAD/NADH balance supports *de novo* Asp biosynthesis and is required for proliferation^37,38^. Since GOT1 is part of the malate-Asp shuttle^37^, we checked whether *GOT1* knockdown affected the NAD/NADH ratio; the NAD/NADH ratio was indeed decreased (**Supplementary Figure 9**), suggesting that this reduction may have influenced the cell proliferation inhibition in both LNCaP and PC3 cells. As depicted in **Supplementary Figure 7**, GOT1 is necessary to convert Asp derived from the Gln TCA cycle into oxaloacetate and malate to produce NADPH^39^, which is essential for maintaining intracellular redox balance via detoxification of damaging reactive oxygen species (ROS). Both LNCaP and PC3 cells showed increased ROS levels upon *GOT1* knockdown (Figure 7e), suggesting that GOT1 plays a role in cellular redox balance and can be manipulated to reduce the viability of prostate cancer cells.

## Discussion

Here we identified a group of putative RNA and metabolite biomarkers in urine and a novel therapeutic target in prostate cancer. To improve the accuracy of disease classification, we carried out metabolic and transcriptomic profiling of urine obtained from BPH, PTT, and PCa patients (without prostatic massage). Urine from normal healthy individuals was used as the control. Through an integrated analysis of metabolomic and transcriptomic data, we identified GOT1 as a key regulator of metabolic changes in PCa patients.

Recent advances in transcriptomics and metabolomics have led to the identification of various candidate biomarkers for cancer diagnosis and prognosis^16,40^. However, biomarkers derived from one dataset may not be reliable, and reproducibility in independent cohorts is challenging^41^. There are great advantages in using biofluids including blood, urine, saliva, and seminal plasma as sources of biomarkers^42,43^. Among them, urine is a promising liquid biopsy as it is noninvasive, replenishable, and convenient to collect. Urine has been at the center of clinical proteomics and provided biomarkers for renal disease^44^, renal cell carcinoma^45^, bladder cancer^46^, and prostate cancer^47^. In addition to proteins and peptides, urine contains various nucleic acids, metabolites, and lipids. Recently, the long noncoding RNA *PCA3* and the fusion gene *TMPRSS2:ERG* have been proposed as urinary PCa biomarkers^48^. Here we report for the first time a global transcriptomic profile of PCa in urine. We applied capture-based enrichment (RNA Access protocol), in which probes target exonic regions, and were able to separate PCa samples from normal healthy individual samples by unsupervised methods (**Supplementary Figure 2**).

In pancreatic ductal adenocarcinoma (PDAC), the transaminase GOT1 is required to sustain cell growth by enabling the production of NADPH to compensate internal ROS (**Supplementary Figure 7**). We showed that GOT1 is essential for prostate cancer cell line (PC3 and LNCaP) growth. GOT1 knockdown increased ROS levels, suggesting that GOT1 may be involved in NADPH generation. As reported previously, GOT1 also functions as a member of the malate-aspartate shuttle^38^, in which two pairs of enzymes, glutamate oxaloacetate transaminases (GOT) and malate dehydrogenase (MDH), serve to transfer reducing equivalents across the mitochondrial membrane (**Supplementary Figure 7**). Our transcriptomic analysis revealed the upregulation of all members of the shuttle including *GOT1*, *GOT2*, *MDH1*, and *MDH2* (**Supplementary Table 7**). These results suggest that the malate-aspartate shuttle may play an important role in cell growth in PCa. This hypothesis is supported by the reduction in NAD/NADH ratio upon GOT1 knockdown in both cell lines.

In conclusion, prostate cancers appear to undergo GOT1-dependent metabolic adaptation to promote a malignant phenotype and resist oxidative stress. The glutamate phenotype represented by the gene expression and metabolic changes in urine reflect this GOT1-dependent pathway in prostate cancer cells. In addition to focusing on these pathway components as biomarkers of prostate cancer in urine, enzymes involved in this pathway might be excellent targets for PCa therapy. Indeed, small molecule inhibitors of GLS1 (mitochondrial glutaminase), which converts glutamine to glutamate, already exist^49^. Targeting this pathway is worthy of further investigation either with or without concurrent ROS-induced cellular stress^49^, this latter approach a particularly appealing strategy in patients with prostate cancers treated with ionizing radiotherapy. Liquid biopsies are an extremely useful tool for non-invasive biomarker and target discovery.

## Materials and Methods

### Sample collection and preparation

Urine samples were collected from 20 benign prostatic hyperplasia (BPH), 11 prostatitis (PTT), 20 prostate cancer (PCa) patients, and 20 normal healthy individuals with no history of cancer at the Global Robotics Institute (Celebration, FL, USA) and Florida Urology Associates (Orlando, FL, USA) of Florida Hospital between 2008-2014. The institutional review board (IRB) of the Florida Hospital approved sample use. Additional 55 urine samples (11 samples from each different gleason score) were obtained from The Johns Hopkins Hospital and the study was approved by IRB of the Johns Hopkins Medical Institutions. All participants were required to sign and provide written consent. Urine samples were collected using urine preservation tubes (Norgen Bioteck, Thorold, ON, Canada) and kept at room temperature until centrifugation to separate the exfoliated cells in the urine samples. Cell-free urine was then stored at −80°C until further use for metabolite analysis. The exfoliated cells from normal and PCa urine samples were used for total RNA purification using the Urine (exfoliated cell) RNA purification kit (Norgen Bioteck). Total RNA was subjected to RNA-seq to identify gene signatures.

### Global untargeted metabolomics

Global metabolomics was performed by ultra-high performance liquid chromatography coupled with high-resolution mass spectrometry (UHPLC-HRMS) on a Thermo Q Exactive with Dionex UHPLC (Thermo Fisher Scientific, Waltham, MA). To 50 μL of urine, 20 μL of internal standard was added (40 μg/mL tryptophan-d3, 4 μg/mL leucine-d10, 4 μg/mL creatine-d3, and 4 μg/mL caffeine-d3) followed by 400 μL of 98:2 acetonitrile:water with 0.1% sodium azide. The solution was vortexed and spun down at 20,000 × g (8C) for 10 min. The supernatant was transferred to a new microcentrifuge tube and dried under a gentle stream of nitrogen. The dried sample was reconstituted in 50 μL of 0.1% formic acid in water and transferred to a LC vial with fused glass insert for analysis. LC-HRMS^50,51^ analysis was performed in positive and negative ion modes as separate injections, injecting 2 μL for positive and 4 μL for negative ions. Separation was achieved on a C18-pfp column (ACE Excel 100×2.1 mm, 2 μm, Advanced Chromatography Technologies, Aberdeen, Scotland) with 0.1% formic acid in water as A and acetonitrile as B. Metabolites were identified by matching to an in-house retention time library of 600 metabolites. All the data normalization, multivariate analyses, pathway analysis, and biomarker discovery were carried out using Metaboanalyst 4.0 (http://www.metaboanalyst.ca). Integrated gene-metabolite network analsyis was conducted using the bioinformatics platform Cytoscape (http://www.cytoscape.org/) with the Metascape plugin (http://metscape.ncibi.org/).

### Cell culture

Prostate cancer cell lines LNCaP (ATCC^®^ CRL-1740™) and PC3 (ATCC^®^ CRL-7934™) were cultured in RPMI 1640 medium and Dulbecco’s Modified Eagle Medium (Thermo Fisher Scientific), respectively, supplemented with 10% FBS and penicillin/streptomycin.

### RNA isolation, cDNA synthesis, and quantitative real-time PCR (qPCR)

Total RNAs from cell lines were purified using the Direct-zol RNA Miniprep kit (Zymo Research, Irvine, CA). Normal prostate epithelial cell RNA was purchased from BioChain Institute Inc. (Catalog # R1234201-50, Newark, CA). RNA (0.5 μg) was then used for cDNA synthesis using a high capacity cDNA reverse transcription kit (Applied Biosystems, Foster City, CA). qPCR was performed using a Power SYBR Green PCR master mix (Applied Biosystems) in the 7500 Real-Time PCR system (Applied Biosystems). A final reaction volume of 10 μl was used containing 1 μl (corresponding to 10 ng) of cDNA template, 5 μl of 2X Power SYBR Green PCR master mix (Applied Biosystems), and 0.2 μM of each primer. The reaction was subjected to denaturation at 95°C for 10 min followed by 40 cycles of denaturation at 95°C for 15 sec and annealing at 58°C for 1 min. SDS1.2.3 software (Applied Biosystems) was used for comparative Ct analysis with TATA-box binding protein (TBP) serving as the endogenous control. The primer sequences for the genes are listed in **Supplementary Table 8**.

### RNA access

The quantity and integrity of the RNA was measured using both the Qubit RNA HS Assay Kit (Thermo Fisher Scientific) and the Agilent 2100 Bioanalyzer RNA Pico kit (Agilent Technologies, Santa Clara, CA). Following the Illumina DV_200_ metric (percentage of RNA fragments greater than 200 nucleotides), 100 ng of RNA with DV_200_ >30% was used to prepare sequencing libraries in accordance with the TruSeq RNA Access protocol (Illumina, Inc., San Diego, CA). First strand cDNA was synthesized using random primers followed by second strand synthesis. The cDNA then underwent 3’ adenylation followed by adapter ligation and PCR amplification (15 cycles). Library quality was measured using both the Qubit dsDNA HS Assay Kit and Agilent Bioanalyzer DNA kit. A 4-plex pool of libraries was then made (200 ng of each sample) followed by two rounds of hybridization/capture and a final amplification (10 cycles). The quality and quantity of the final libraries were determined using the Agilent 2100 Bioanalyzer DNA HS kit and Kapa Biosystems qPCR (Kapa Biosystems, Inc., Wilmington, MA). Multiplexed libraries were pooled and normalized to 17.5 pM. The libraries were sequenced using a 75 bp paired-end run on the Illumina MiSeq instrument. Paired-end reads were mapped to the human genome (hg19) using tophat2.0.1; mapped reads were filtered based on the mapping quality. The overall mapping rates were about 93%. mRNA quantification was conducted in Partek Genomics Suite 6.6. R package edgeR was used to analyze the differential expression of mRNAs.

### Clustering and principal component analysis (PCA)

The resulting mRNA expression profile included 46,459 transcripts, and the non-parametric Mann-Whitney U-test was used to identify significantly regulated transcripts. 5510 transcripts were identified (p≤0.05) as significantly differentially expressed between normal and PCa groups. Within those 5,510 transcripts, 1,118 transcripts had RPKM values greater than 1.0 for all samples. We pre-compiled a gene panel that lists all prostate cancer-related genes (with the help of Illumina). By comparing with this panel, we obtained 542 transcripts with RPKMs greater than 1.0 for all samples. Within these 542, 116 transcripts were significantly regulated (Table 1). All overlapping transcripts between the expression profile and prostate cancer panel, a total of 3825 transcripts, were used to run the unsupervised clustering analysis and PCA. Hierarchical cluster analysis was performed in R using the correlation between samples to characterize similarity. Initially, each sample was assigned to its own cluster and then the algorithm proceeded iteratively, at each stage joining the two most similar clusters and continuing until there was just a single cluster. Correlation between samples was calculated using the expression values of the 3825 transcripts. We also used PCA to visualize sample to sample distance. The transformation was defined that the first principal component accounted for the largest variance (as much of the variability in the dataset as possible). In the results, each sample was projected onto the 3D space in which the three axes were the first three highest principle components (see **Supplementary Figure 2b**).

### Gene set enrichment analysis (GSEA) of tissue RNA-seq data

Gene expression was analyzed in 65 patients using RNA-seq data^23^. Briefly, RPKM values from tumor and matched normal samples from 65 patients were analyzed using paired linear model differential expression analysis using the Bioconductor limma package^52^ to find differentially expressed genes between tumor and matched normal. To compare between groups for tumor or matched normal tissues, linear model analysis using the limma package was used on RPKM values for all genes. The analysis was performed in R (version 3.4.4, www.R-project.org).

GSEA for KEGG pathways was conducted using the GSEA desktop application (borad.mit.edu/gsea). All genes were ranked using scores based on fold-change direction and p-value, and enrichment analysis was conducted using GSEAPreranked with the ‘classic’ enrichment statistic. Significantly enriched genes sets were identified using q-value ≤0.05 as a cutoff.

### Transient transfection and cell proliferation assay

Cells (0.3 × 10^6^ cells) were mixed with siRNA (Thermo Fisher Scientific, final concentration 20 nM) and lipofectamine RNAiMAX (Thermo Fisher Scientific) mixture in 2 ml medium containing 10% FBS. Cells were plated in duplicate at 7500 cells per well into 96-well plates and, after 24 h, the medium was replaced. Cell proliferation was assessed using the CellTiter96 Aqueous One Solution Cell Proliferation Assay (MTS) kit (Promega, Madison, WI).

### NAD and NADH quantification

NAD and NADH levels were measured using the NAD/NADH-Glo Assay kit (Promega). Cells were plated in duplicate (7500 cells per well) into 96-well plates. Cells were lysed with base solution (100 mM sodium carbonate, 20 mM sodium bicarbonate, 10 mM nicotinamide, and 0.05% Triton X-100) with 1% DTAB (Sigma, D8638). Lysates were heat-treated at 60°C for 20 min in the presence/absence of acid. Heat-treated samples were then subjected to the luciferase assay according to the manufacturer’s protocol and the NAD/NADH ratio was calculated.

### Measurement of reactive oxygen species (ROS)

Cells (1 × 10^6^ cells) were resuspended in 1 ml medium containing 20 uM 2’7’-dichlorofluoresicin diacetate (DCFH-DA) (Sigma, D6883) and incubated for 30 min at 37°C in 5% CO_2_. Fluorescent cells were detected using a FACSCalibur flow cytometer (Becton Dickinson, Anaheim, CA) and data were analyzed with the Software 2.5.1 from Flowing Software (www.flowingsoftware.com).

### Invasion assay

The cell invasion assay was performed using Corning BioCoat Matrigel Invasion Chambers (Discovery Labware, Bedford, MA) according to the manufacturer’s protocol. Briefly, cells were starved in serum-free medium for 24 h and plated into the upper chambers in serum-free medium (0.2 × 10^6^ cells per chamber). Medium containing 10% FBS was added to the lower chambers. Cells were incubated for 48 h at 37°C. Invaded cells were stained with 0.5% crystal violet dye. After washing the excess dye, cells were air dried. Methanol was used to extract dye from cells and optical density was measured at 570 nm.

### Soft agar colony formation assay

To access anchorage-independent growth, a CytoSelect 96-well cell transformation assay kit (Cell Biolabs Inc., San Diego, CA) was used according to the manufacturer’s protocol. Briefly, cells were seeded in soft agar at a density of 10,000 cells per well and incubated at 37°C in 5% CO_2_ for 7 days. Colony formation was quantified by the MTT assay provided with the kit according to the assay protocol.

## Acknowledgments

We thank Sanford Burnham Prebys Medical Discovery Institute (SBP) Analytical Genomics core facility for deep-sequencing, Bioinformatics core for data analysis support, and Ms. Debbie McFadden for formatting the manuscript. The authors thank Drs. Andrei Osterman and David Scott at SBP for comments and suggestions.

## Conflicts of interest

There are no conflicts of interest.

