## Supplementary table and figures for "Integrated RNA and metabolite profiling of urine liquid biopsies for prostate cancer biomarker discovery"

**Supplementary Information**

**Supplementary Table 1.** The clinicopathological details of patients with prostate cancer providing urine for RNA-seq.

**Supplementary Table 2.** The expression of known prostate cancer marker genes in prostate cancer urine samples.

**Supplementary Table 3.** Pathways represented by genes differentially expressed in the cells extracted from the urine of normal and cancer patients.

**Supplementary Table 4.** A comparison of expression changes of genes differentially expressed in the urine of patients with prostate cancer compared to normal in The Cancer Genome Atlas data.

**Supplementary Table 5.** A comparison of urine RNA access-seq with prostate tissue RNA-seq.

**Supplementary Table 6.** The top ten pathways most significantly enriched for differentially expressed genes and metabolites using an integrated analysis in MetaboAnalyst 3.0.

**Supplementary Table 7.** The expression of TCA cycle and glutamine metabolism genes in prostate cancer urine samples.

**Supplementary Table 8.** qPCR primer sequences.

**Supplementary Figure 1.** Electropherograms of RNA extracted from (a) LNCaP and PC3 prostate cancer cell lines and normal prostate tissue (control); (b) RNA from the urine of “normal” patients; and (c) RNA from the urine of patients with cancer. RNA extracted from urine is generally severely degraded.

**Supplementary Figure 2.** Unsupervised clustering [using Treeview, a) and principal component analysis (PCA), b)] of significantly differentially expressed genes detected in the cells extracted from urine of normal (green) and cancer (red) patients. Normal and cancer specimens are readily but not perfectly separated.

**Supplementary Figure 3.** The identified subgroups do not differ with respect to clinical features. The bars represent the percentage of different categories of (A) Gleason score (3+3, 3+4, 4+3 and ≥4+4); (B) tumor stages (T2, T3, T4); and (C) metastasis (localized, localized advanced, metastasized) found in groups A and B. Fisher’s exact test showed no significant difference of distribution of patients for any clinical trait within the two groups.

**Supplementary Figure 4.** PCA3 and KLK3 expression in groups A and B.

**Supplementary Figure 5.** Global untargeted urine metabolomics data normalization and metabolite profile analysis. (A) Box plots and kernel density plots before and after normalization (Auto Scaling). The box plots show at most 40 metabolic features and the density plots are based on all samples. The graph summarizes the distribution of input data values before and after normalization. The box plots on the top show the concentration distributions of individual metabolic features, whereas the bottom plots show the overall concentration distribution based on kernel density (Auto Scaling) estimation.  (B) Multivariate principal component analysis (PCA) scores plot of repeated samples for normal, BPH (benign prostate hyperplasia), PTT (prostatitis), and PCa (prostate cancer). (C) Unbiased hierarchical clustering-based heatmap analysis shows global untargeted metabolite profile for normal, BPH, PTT, and PCa urine samples. Red indicates the increasing direction, and blue indicates the decreasing direction. (C) Metabolic pathway analysis plot created using MetaboAnalyst 4.0. Plots depict different metabolic pathways that are increased in PCa urine samples compared to normal. The x-axis represents the pathway impact value computed from pathway topological analysis, and the y-axis is the -log of the P-value obtained from pathway enrichment analysis. The pathways that were most significantly changed are characterized by both a high -log(p) value and high impact value (top right region). The following are the top 14 metabolic pathways highly upregulated in PCa urine:

1. Alanine, aspartate and glutamate metabolism
2. Citrate cycle (TCA cycle)
3. Pyruvate metabolism
4. Valine, leucine and isoleucine degradation
5. D-Glutamine and D-glutamate metabolism
6. Butanoate metabolism
7. Propanoate metabolism
8. Glyoxylate and dicarboxylate metabolism
9. Arginine and proline metabolism
10. Glycolysis / Gluconeogenesis
11. Valine, leucine and isoleucine biosynthesis
12. Synthesis and degradation of ketone bodies
13. Tryptophan metabolism
14. Phenylalanine, tyrosine and tryptophan biosynthesis

**Supplementary Figure 6.** Gene-metabolite network in PCa urine samples. Joint omics analysis revealed alanine, aspartate and glutamate metabolism, citrate cycle (TCA cycle), and D-glutamine and D-glutamate metabolism pathway metabolites as highly enriched in PCa urine samples compared to normal.

**Supplementary Figure 7.** Schematic view of TCA cycle and glutamine metabolism. Red arrows indicate up-regulated genes in PCa urine compared to normal urine samples.

**Supplementary Figure 8.** Glutamate and Aspartate level changes in *GOT1* knockdown LNCaP and PC3 prostate cancer cell lines. The data from three independent experiments were expressed as mean ± SD.

**Supplementary Figure 9.** NAD/NADH ratios in LNCaP and PC3 prostate cancer cell lines. The data from two independent experiments were expressed as mean ± SD.

**Supplementary Table 1.** The clinicopathological details of patients with prostate cancer providing urine for RNA-seq.

| **Sample ID** | **Gender** | **PSA** | **Pathological Stage** | **Gleason grade** | **Age at diagnosis** | **Diagnosis** | **LN Positive** |
| --- | --- | --- | --- | --- | --- | --- | --- |
| 1C | M | 2.7 | pT2a | 6 | 48 | Prostatic Adenocarcinoma | Not submitted |
| 2C | M |  |  |  |  |  |  |
| 3C | M | 5.5 | pT2c | 6 | 58 | Prostatic Adenocarcinoma | Negative for tumor |
| 4C | M | 4.7 | pT2c | 7 | 69 | Prostatic Adenocarcinoma | Negative for tumor |
| 5C | M | 5.2 | pT3a | 7 | 69 | Prostatic Adenocarcinoma | Not submitted |
| 6C | M | 3.5 | pT2c | N/A |  | Prostatic Adenocarcinoma | Negative for tumor |
| 7C | M | 2.2 | pT3a | 8 | 67 | Prostatic Adenocarcinoma | Metastatic |
| 9C | M | 11.8 | pT3a | 7 | 73 | Prostatic Adenocarcinoma | Not submitted |
| 8C | M | 7.9 | pT2c | 6 | 61 | Prostatic Adenocarcinoma | Not submitted |
| 10C | M | 4.2 | pT2a | 6 | 57 | Prostatic Adenocarcinoma | Negative for tumor |
| 11C | M | 4.6 | pT2c | 7 | 71 | Prostatic Adenocarcinoma | Not submitted |

M: male, PSA: prostate-specific antigen level (ng/ml), LN: lymph node, N/A: not available

**Supplementary Table 2**. The expression of known prostate cancer marker genes in prostate cancer urine samples.

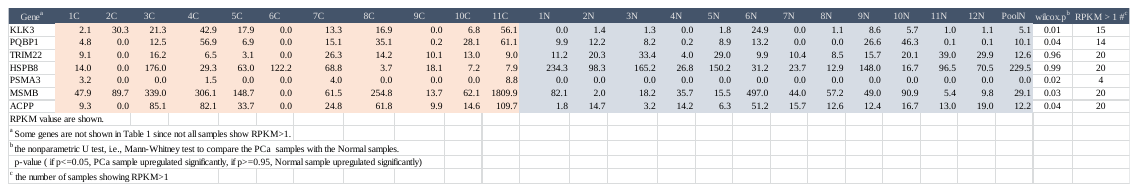

**Supplementary Table 3.** Pathways represented by genes differentially expressed in the cells extracted from the urine of normal and cancer patients.

| **Ingenuity Canonical Pathways** | **-Log (p-value)** | **Genes** |
| --- | --- | --- |
| Molecular Mechanisms of Cancer | 10.5 | *RAF1,GNAS,SMAD3,PIK3R1,CREBBP,TYK2,GNAQ,NCSTN,MDM2,JAK2,XIAP,EP300,PTK2,SMAD4,CDKN1B,NOTCH1,BIRC3* |
| Pancreatic Adenocarcinoma Signaling | 7.61 | *RAF1,PIK3R1,SMAD3,TYK2,SMAD4,MDM2,JAK2,CDKN1B,NOTCH1* |
| PPARα/RXRα Activation | 7.42 | *RAF1,GNAS,HSP90AB1,PLCG2,SMAD3,CREBBP,GNAQ,SMAD4,JAK2,EP300* |
| Chronic Myeloid Leukemia Signaling | 6.88 | *HDAC6,RAF1,HDAC4,PIK3R1,SMAD3,SMAD4,MDM2,CDKN1B* |
| IL-15 Production | 6.39 | *PTK2,PTK2B,TYK2,JAK2,IRF1* |
| Renin-Angiotensin Signaling | 6.39 | *PTK2,RAF1,PTK2B,PLCG2,PIK3R1,GNAQ,JAK2,ACE* |
| Prolactin Signaling | 6.32 | *RAF1,PLCG2,PIK3R1,CREBBP,JAK2,IRF1,EP300* |
| PI3K/AKT Signaling | 6.31 | *RAF1,HSP90AB1,PIK3R1,TYK2,MDM2,JAK2,CDKN1B,MCL1* |
| NF-κB Signaling | 6.1 | *RAF1,MYD88,BCL10,PLCG2,PIK3R1,CREBBP,IGF1R,TNFAIP3,EP300* |
| Prostate Cancer Signaling | 6.02 | *RAF1,HSP90AB1,PIK3R1,CREBBP,MDM2,CDKN1B,EP300* |
| Hereditary Breast Cancer Signaling | 5.88 | *HDAC6,NPM1,HDAC4,PIK3R1,SMARCB1,CREBBP,DDB2,EP300* |
| Role of NFAT in Cardiac Hypertrophy | 5.86 | *HDAC6,RAF1,GNAS,HDAC4,PLCG2,PIK3R1,IGF1R,GNAQ,EP300* |
| Protein Kinase A Signaling | 5.82 | *PTK2,RAF1,GNAS,PTPN2,ATF1,PTK2B,PLCG2,SMAD3,CREBBP,GNAQ,SMAD4,EP300* |
| Cell Cycle: G1/S Checkpoint Regulation | 5.8 | *HDAC6,HDAC4,SMAD3,SMAD4,MDM2,CDKN1B* |
| ERK/MAPK Signaling | 5.69 | *PTK2,RAF1,ELF4,ATF1,PTK2B,PLCG2,PIK3R1,CREBBP,EP300* |
| Mouse Embryonic Stem Cell Pluripotency | 5.63 | *RAF1,PIK3R1,TYK2,CREBBP,SMAD4,JAK2,XIAP* |
| Telomerase Signaling | 5.55 | *HDAC6,TERF2,RAF1,ELF4,HDAC4,HSP90AB1,PIK3R1* |
| iNOS Signaling | 5.35 | *MYD88,TYK2,CREBBP,JAK2,IRF1* |
| Glucocorticoid Receptor Signaling | 5.28 | *RAF1,ICAM1,HSP90AB1,PIK3R1,SMAD3,SMARCB1,CREBBP,SMAD4,JAK2,EP300* |
| Tec Kinase Signaling | 5.26 | *PTK2,GNAS,PTK2B,PLCG2,PIK3R1,TYK2,GNAQ,JAK2* |

**Supplementary Table 4.** A comparison of expression changes of genes differentially expressed in the urine of patients with prostate cancer compared to normal in The Cancer Genome Atlas data.

| **Gene** | **Normal**  **Mean log2 (RSEM+1)** | **SD** | **Tumor**  **Mean log2 (RSEM+1)** | **SD** | **Δ mean log2 (RSEM+1)** | **p-value** | **Up / downregulated** |
| --- | --- | --- | --- | --- | --- | --- | --- |
| *ACE* | 10.2 | 1.2 | 9.1 | 0.9 | -1.1 | <0.0001 | Down |
| *ATF1* | 8.9 | 0.3 | 8.7 | 0.5 | -0.2 | 0.0011 | Down |
| *BRD3* | 10.4 | 0.3 | 10.5 | 0.5 | 0.2 | 0.0094 | Up |
| *CCNB1IP1* | 9.3 | 0.4 | 9.1 | 0.5 | -0.2 | 0.0084 | Down |
| *CDC14A* | 8.5 | 0.9 | 8.7 | 0.9 | 0.2 | 0.0780 |  |
| *CDK8* | 7.6 | 0.4 | 7.1 | 0.7 | -0.5 | 0.0000 | Down |
| *ELK4* | 8.2 | 0.9 | 8.5 | 0.8 | 0.2 | 0.0471 | Up |
| *EPCAM* | 11.3 | 0.8 | 12.5 | 0.7 | 1.2 | <0.0001 | Up |
| *FH* | 10.2 | 0.3 | 10.4 | 0.3 | 0.2 | 0.0006 | Up |
| *GMPS* | 9.6 | 0.3 | 9.6 | 0.4 | 0.1 | 0.2340 |  |
| *GNAS* | 14.3 | 0.3 | 14.3 | 0.4 | 0.0 | 0.5377 |  |
| *GOT1* | 10.0 | 0.6 | 9.8 | 0.4 | -0.2 | 0.0025 | Down |
| *GRHPR* | 10.9 | 0.3 | 11.2 | 0.3 | 0.3 | <0.0001 | Up |
| *HDAC6* | 9.9 | 0.4 | 10.0 | 0.3 | 0.1 | 0.0358 | Up |
| *HSP90AB1* | 14.7 | 0.3 | 14.9 | 0.7 | 0.2 | 0.0659 |  |
| *LRPPRC* | 11.4 | 0.2 | 11.5 | 0.4 | 0.1 | 0.0944 |  |
| *MSH3* | 8.9 | 0.3 | 8.8 | 0.5 | -0.1 | 0.3042 |  |
| *NACA* | 13.8 | 0.4 | 14.2 | 0.5 | 0.4 | <0.0001 | Up |
| *NPM1* | 12.8 | 0.4 | 13.3 | 0.4 | 0.5 | <0.0001 | Up |
| *PFDN5* | 12.3 | 0.5 | 12.3 | 0.6 | 0.0 | 0.7540 |  |
| *PHB* | 11.5 | 0.3 | 11.6 | 0.4 | 0.1 | 0.0224 | Up |
| *PHF6* | 9.3 | 0.5 | 9.3 | 0.6 | -0.1 | 0.3891 |  |
| *PIK3R1* | 11.4 | 0.6 | 10.3 | 0.7 | -1.1 | <0.0001 | Down |
| *PTK2* | 11.1 | 0.2 | 11.0 | 0.4 | -0.1 | 0.1724 |  |
| *PTPN2* | 8.8 | 0.3 | 8.8 | 0.3 | 0.0 | 0.4067 |  |
| *RPL22* | 12.9 | 0.3 | 13.2 | 0.4 | 0.3 | <0.0001 | Up |
| *RPS11* | 14.6 | 0.5 | 15.0 | 0.6 | 0.4 | <0.0001 | Up |
| *SDHC* | 10.8 | 0.2 | 10.8 | 0.3 | 0.0 | 0.4445 |  |
| *SDHD* | 11.0 | 0.3 | 10.4 | 0.4 | -0.6 | <0.0001 | Down |
| *SMAD4* | 11.4 | 0.3 | 11.1 | 0.4 | -0.3 | <0.0001 | Down |
| *SMARCB1* | 10.6 | 0.3 | 10.7 | 0.3 | 0.1 | 0.0059 | Up |
| *TCEA1* | 10.6 | 0.3 | 10.6 | 0.4 | 0.0 | 0.6120 |  |
| *TERF2* | 9.2 | 0.2 | 9.2 | 0.4 | -0.1 | 0.1909 |  |
| *TFG* | 10.9 | 0.5 | 11.0 | 0.3 | 0.1 | 0.0114 | Up |
| *TMEM230* | 11.6 | 0.4 | 11.6 | 0.3 | -0.1 | 0.1750 |  |
| *ZMYND11* | 11.6 | 0.3 | 11.1 | 0.4 | -0.4 | <0.0001 | Down |
| *ZNF585B* | 6.4 | 0.5 | 5.9 | 0.8 | -0.5 | <0.0001 | Down |

**Supplementary Table 5.** A comparison of urine RNA access-seq with prostate tissue RNA-seq.

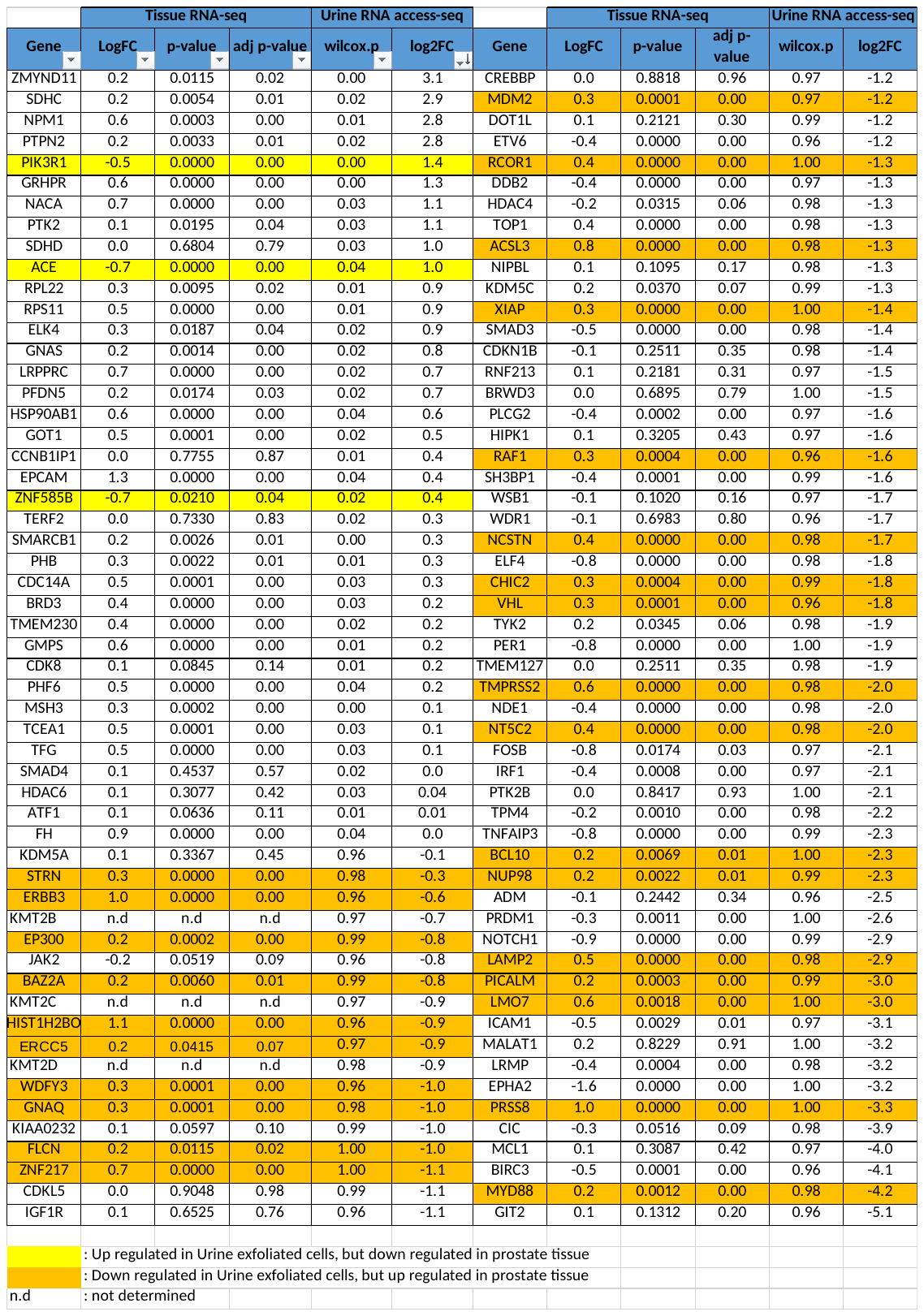

**Supplementary Table 6.** The top ten pathways most significantly enriched for differentially expressed genes and metabolites using an integrated analysis in MetaboAnalyst 3.0.

| **Pathway** | **Total** | **Expected** | **Hits** | **p-value** | **Topology** |
| --- | --- | --- | --- | --- | --- |
| Aminoacyl-tRNA biosynthesis | 87 | 0.85134 | 10 | 2.912E-9 | 0.14493 |
| Alanine, aspartate and glutamate metabolism | 56 | 0.54799 | 6 | 1.1105E-5 | 0.55102 |
| Citrate cycle (TCA cycle) | 50 | 0.48927 | 5 | 9.4033E-5 | 0.36364 |
| Arginine and proline metabolism | 102 | 0.99812 | 6 | 3.4034E-4 | 0.21505 |
| D-Glutamine and D-glutamate metabolism | 9 | 0.088069 | 2 | 0.0031787 | 0.28571 |
| Phenylalanine, tyrosine and tryptophan biosynthesis | 9 | 0.088069 | 2 | 0.0031787 | 1.4 |
| Valine, leucine and isoleucine biosynthesis | 13 | 0.12721 | 2 | 0.0067233 | 0.18182 |
| Histidine metabolism | 44 | 0.43056 | 3 | 0.0084469 | 0.3125 |
| Cysteine and methionine metabolism | 63 | 0.61648 | 3 | 0.022399 | 0.32727 |
| Phenylalanine metabolism | 29 | 0.28378 | 2 | 0.031798 | 0.22727 |

**Supplementary Table 7.** The expression of TCA cycle and glutamine metabolism genes in prostate cancer urine samples.

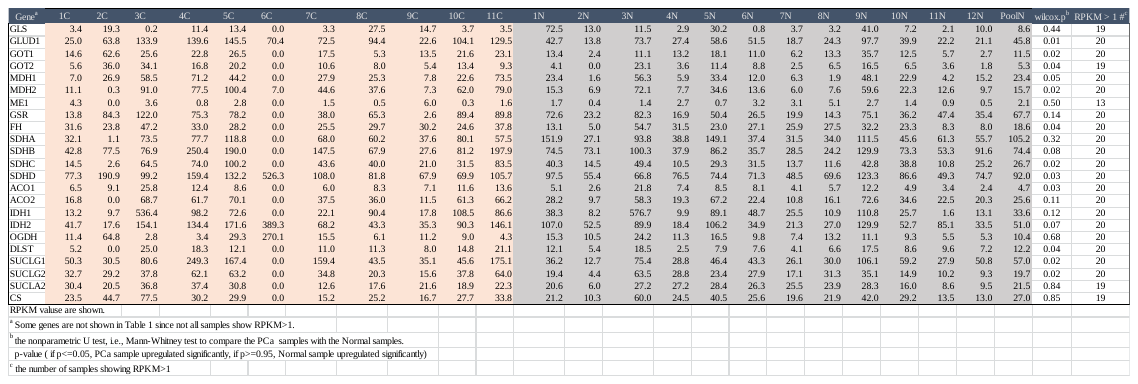

**Supplementary Table 8.** qPCR primer sequences.

| **Oligos** | **Sequence** | **Amplicon size (bp)** |
| --- | --- | --- |
| BRD3 qPCR F | CTGAAACCCACCACTTTGCG | 84 |
| BRD3 qPCR R | GCTTGCTGAGAACGGTTTCC |  |
| ELK4 qPCR F | GCCCCTTGCTCTCCAGTATC | 147 |
| ELK4 qPCR R | CATCCAGCCCAGACAGAGTG |  |
| EPCAM qPCR F | GTTCGGGCTTCTGCTTGC | 89 |
| EPCAM qPCR R | CAGTTTACGGCCAGCTTGTA |  |
| FH qPCR F | CCGAGCACTTCGGCTCCT | 150 |
| FH qPCR R | ATCCGGAAGGAATTTTGGCTTG |  |
| GRHPR qPCR F2 | GACCACGTGGACAAGAGGAT | 146 |
| GRHPR qPCR R2 | TCTGTCAGGACATCTGGGGT |  |
| HDAC6 qPCR F | GCTGACTACCTAGCTGCCTG | 89 |
| HDAC6 qPCR R | TCAAAGCCAGCTGAGACCAG |  |
| NACA qPCR F | CACGCTCTCGCTCGGTCTTT | 148 |
| NACA qPCR R | GGCTGAAGACATAGGTAGCACA |  |
| NPM1 qPCR F | ACTCCAGCCAAAAATGCACA | 208 |
| NPM1 qPCR R | TACATGTAGTGCCCAGGACTGTT |  |
| PHB qPCR F | ATCACCCAGAGAGAGCTGGT | 132 |
| PHB qPCR R | CACCGCTTCTGTGAACTCCT |  |
| RPL22 qPCR F | TGACATCCGAGGTGCCTTTC | 101 |
| RPL22 qPCR R | GTTAGCAACTACGCGCAACC |  |
| RPS11 qPCR F | GTACACCTGTCCCCCTGCTT | 91 |
| RPS11 qPCR R | TGAAGCGCACTGTCTTGCT |  |
| SMARCB1 qPCR F2 | GCGAGTTCTACATGATCGGCT | 145 |
| SMARCB1 qPCR R2 | CCGTGATCATGTGACGATGC |  |
| TFG qPCR F | ATCGTTCAGGAACACCCGAC | 147 |
| TFG qPCR R | CCTGTTGCTGGTACTGTTGG |  |
| GOT1 qPCR F | AGAAGCCCTCAAAACCCCTG | 143 |
| GOT1 qPCR R | CGTTGATTCGACCACTTGGC |  |
| HPRT1 E2 qPCR F | TGCTGAGGATTTGGAAAGGGT | 99 |
| HPRT1 E3 qPCR R | TGATGGCCTCCCATCTCCTT |  |
| TBP E2 qPCR F | ACAACAGCCTGCCACCTTAC | 96 |
| TBP E3 qPCR R | TGCCATAAGGCATCATTGGACT |  |
| ACTB qPCR F | CCTGGCATTGCCGACAGGATG | 107 |
| ACTB qPCR R | CCGATCCACACGGAGTACTTGCG |  |

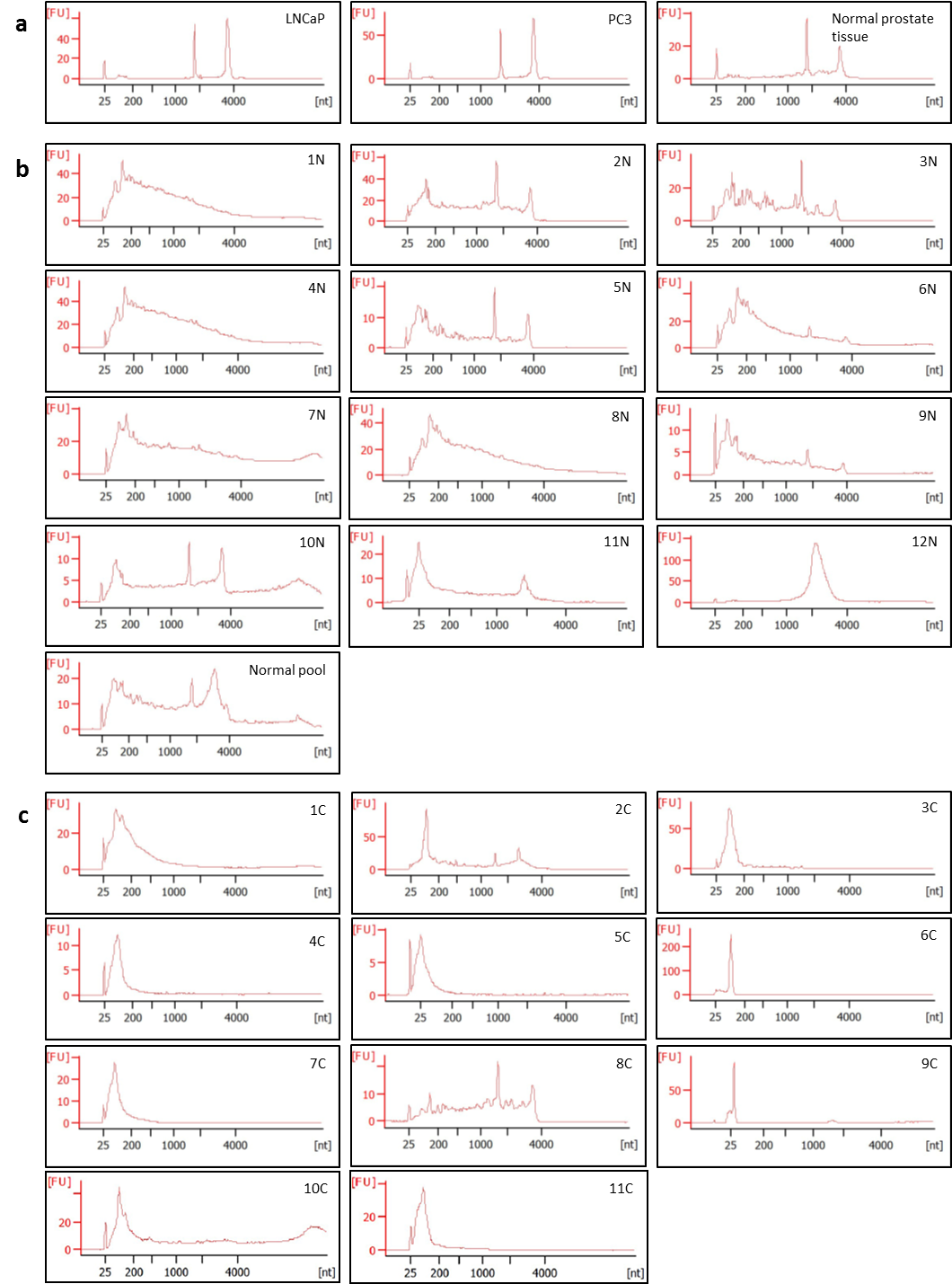

**Supplementary Figure 1.** Electropherograms of RNA extracted from (a) LNCaP and PC3 prostate cancer cell lines and normal prostate tissue (control); (b) RNA from the urine of “normal” patients; and (c) RNA from the urine of patients with cancer. RNA extracted from urine is generally severely degraded.

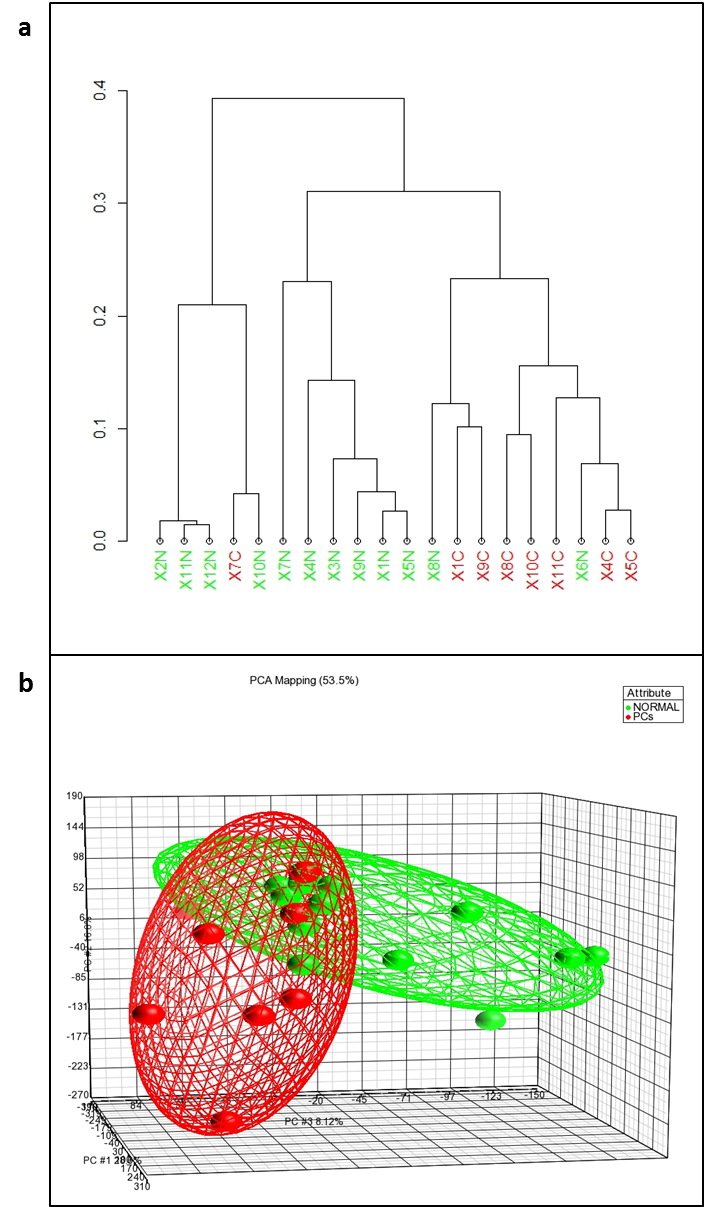

**Supplementary Figure 2.** Unsupervised clustering [using Treeview, a) and principal component analysis (PCA), b)] of significantly differentially expressed genes detected in the cells extracted from urine of normal (green) and cancer (red) patients. Normal and cancer specimens are readily but not perfectly separated.

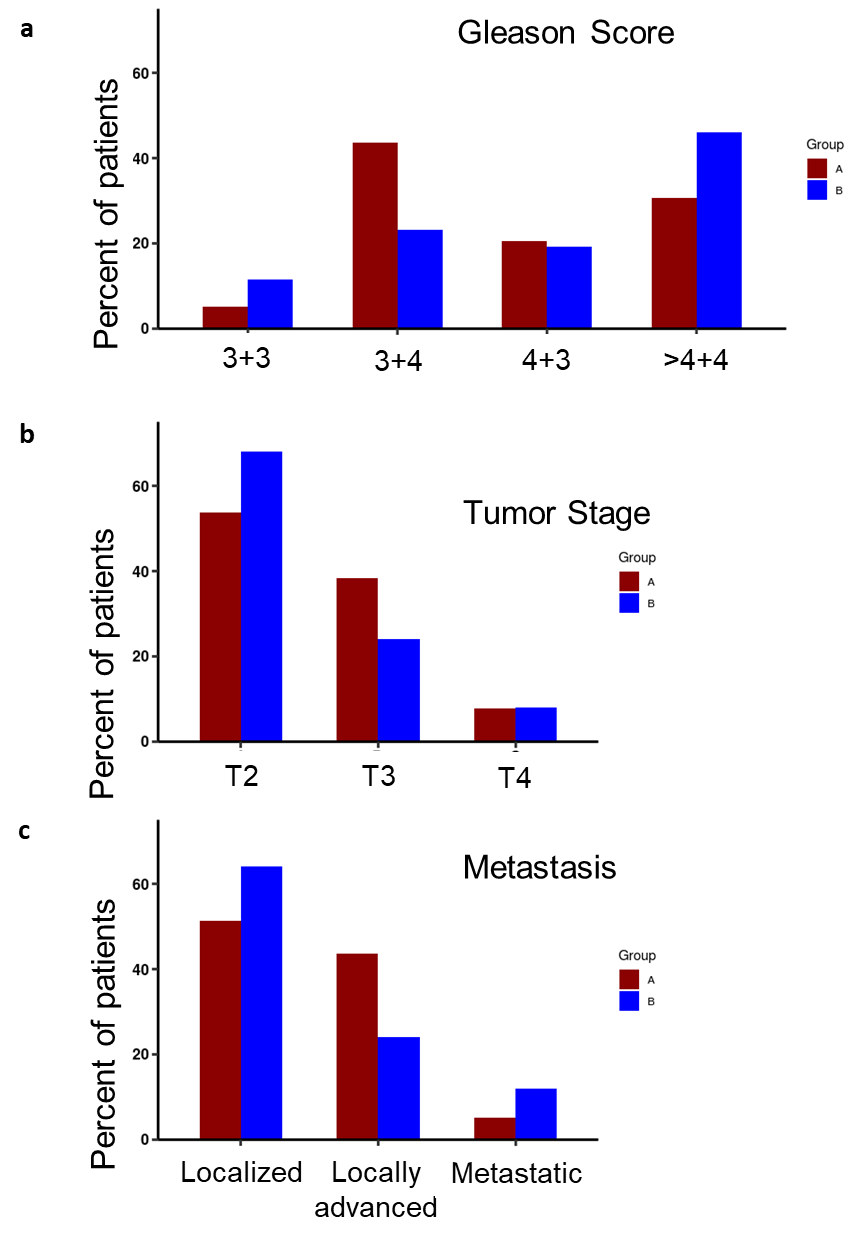

**Supplementary Figure 3.** The identified subgroups do not differ with respect to clinical features. The bars represent the percentage of different categories of (A) Gleason score (3+3, 3+4, 4+3 and ≥4+4); (B) tumor stages (T2, T3, T4); and (C) metastasis (localized, localized advanced, metastasized) found in groups A and B. Fisher’s exact test showed no significant difference of distribution of patients for any clinical trait within the two groups.

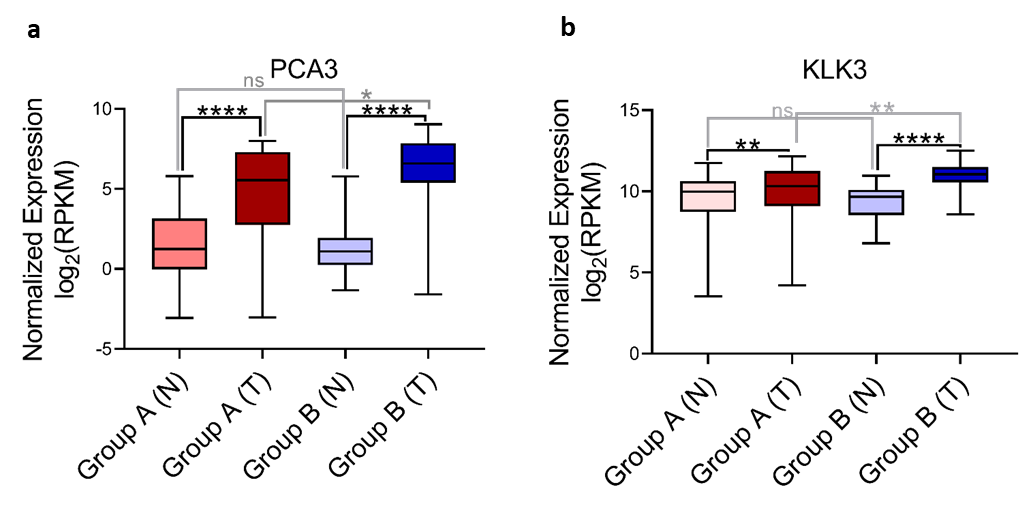

**Supplementary Figure 4.** *PCA3* and *KLK3* expression in groups A and B.

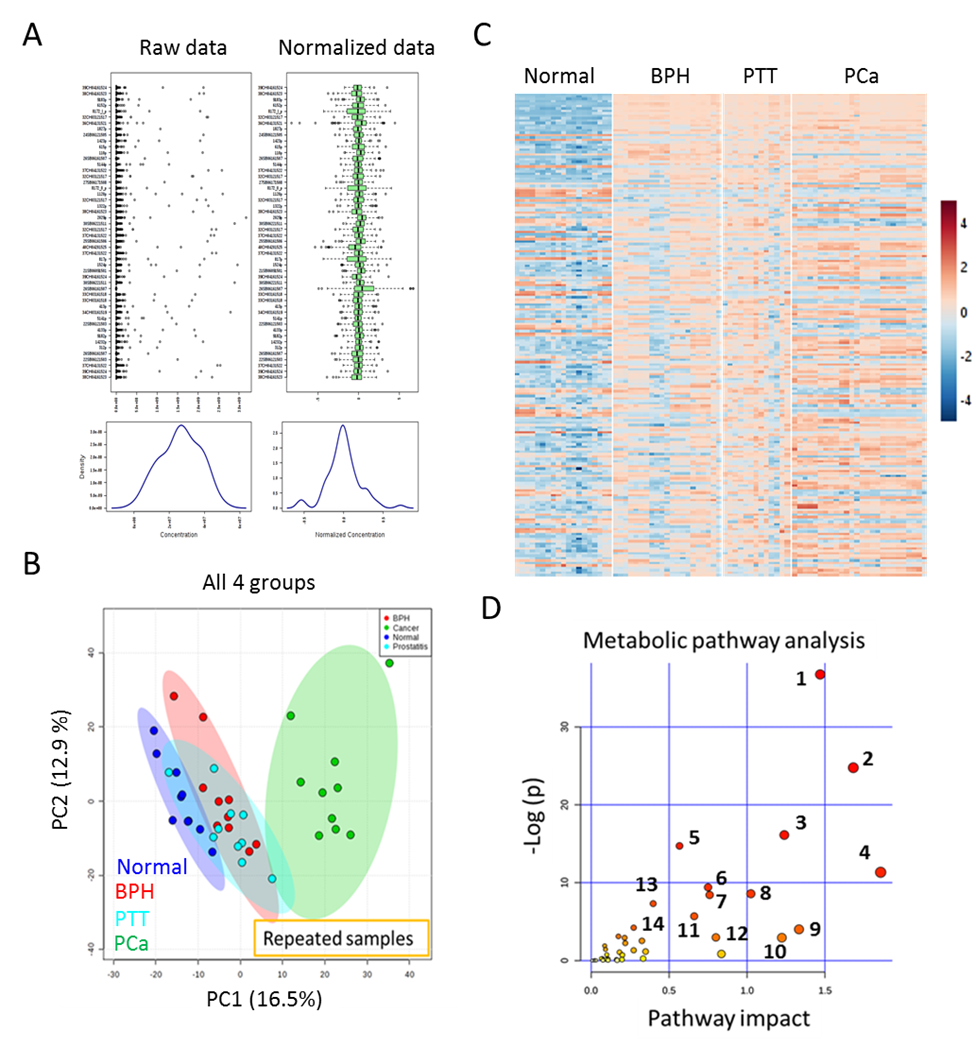

**Supplementary Figure 5.** Global untargeted urine metabolomics data normalization and metabolite profile analysis. (A) Box plots and kernel density plots before and after normalization (Auto Scaling). The box plots show at most 40 metabolic features and the density plots are based on all samples. The graph summarizes the distribution of input data values before and after normalization. The box plots on the top show the concentration distributions of individual metabolic features, whereas the bottom plots show the overall concentration distribution based on kernel density (Auto Scaling) estimation.  (B) Multivariate principal component analysis (PCA) scores plot of repeated samples for normal, BPH (benign prostate hyperplasia), PTT (prostatitis), and PCa (prostate cancer). (C) Unbiased hierarchical clustering-based heatmap analysis shows global untargeted metabolite profile for normal, BPH, PTT, and PCa urine samples. Red indicates the increasing direction, and blue indicates the decreasing direction. (C) Metabolic pathway analysis plot created using MetaboAnalyst 4.0. Plots depict different metabolic pathways that are increased in PCa urine samples compared to normal. The x-axis represents the pathway impact value computed from pathway topological analysis, and the y-axis is the -log of the P-value obtained from pathway enrichment analysis. The pathways that were most significantly changed are characterized by both a high -log(p) value and high impact value (top right region). The following are the top 14 metabolic pathways highly upregulated in PCa urine:

1. Alanine, aspartate and glutamate metabolism
2. Citrate cycle (TCA cycle)
3. Pyruvate metabolism
4. Valine, leucine and isoleucine degradation
5. D-Glutamine and D-glutamate metabolism
6. Butanoate metabolism
7. Propanoate metabolism
8. Glyoxylate and dicarboxylate metabolism
9. Arginine and proline metabolism
10. Glycolysis / Gluconeogenesis
11. Valine, leucine and isoleucine biosynthesis
12. Synthesis and degradation of ketone bodies
13. Tryptophan metabolism

14. Phenylalanine, tyrosine and tryptophan biosynthesis

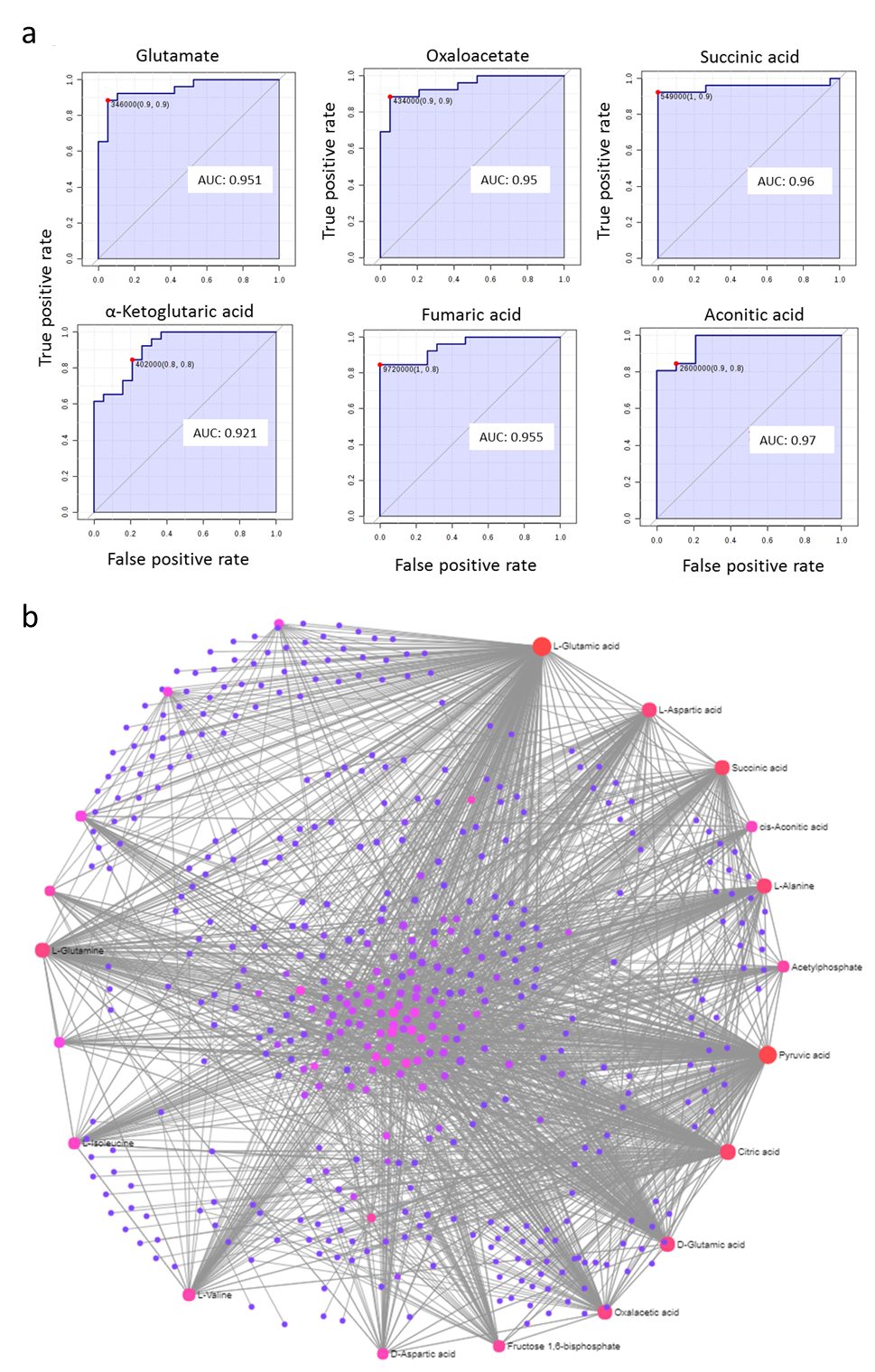

**Supplementary Figure 6.** a) Top six qualified metabolite-based urine biomarkers (aconitic acid, succinic acid, glutamate, oxaloacetate, fumaric acid, and α-ketoglutaric acid) for PCa diagnosis. AUC, area under the ROC curve. b) Gene-metabolite network in PCa urine samples. Joint omics analysis revealed alanine, aspartate and glutamate metabolism, citrate cycle (TCA cycle), and D-glutamine and D-glutamate metabolism pathway metabolites as highly enriched in PCa urine samples compared to normal.

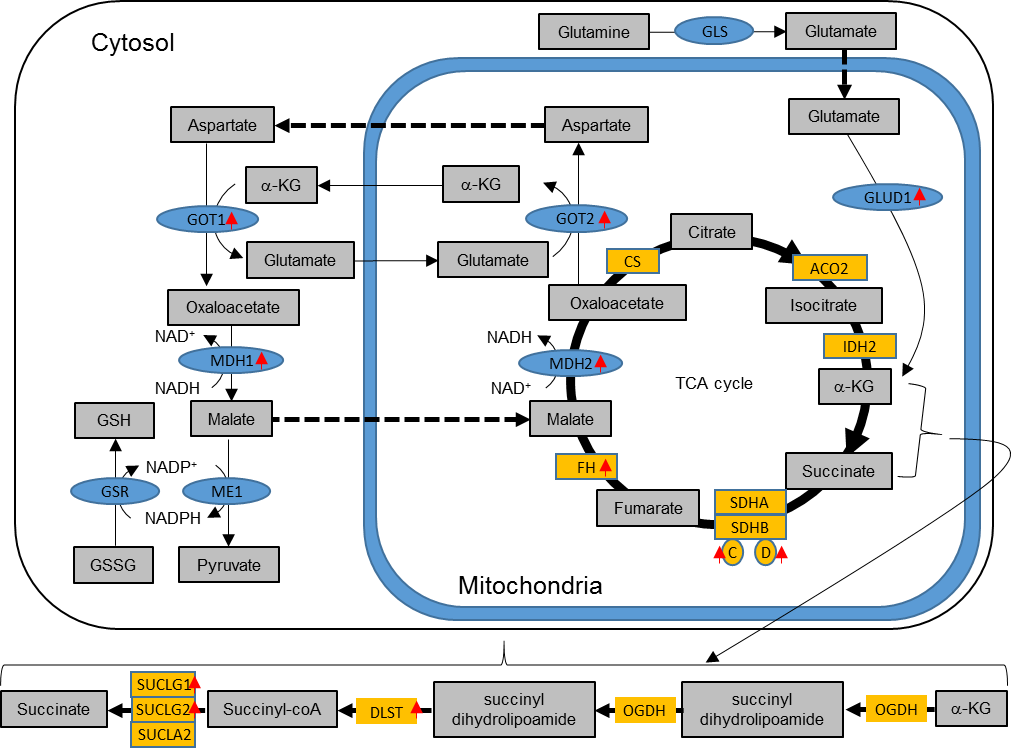

**Supplementary Figure 7.** Schematic view of TCA cycle and glutamine metabolism. Red arrows indicate upregulated genes in PCa urine compared to normal urine samples.

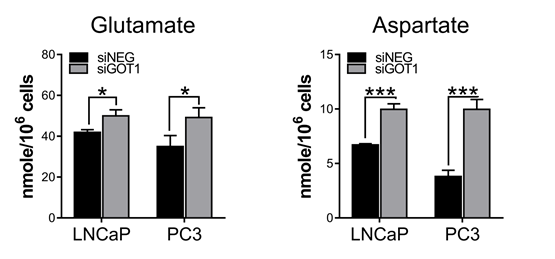

**Supplementary Figure 8.** Glutamate and Aspartate level changes in *GOT1* knockdown LNCaP and PC3 prostate cancer cell lines. The data from three independent experiments were expressed as mean ± SD.

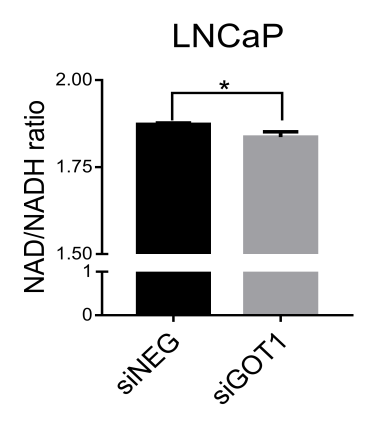

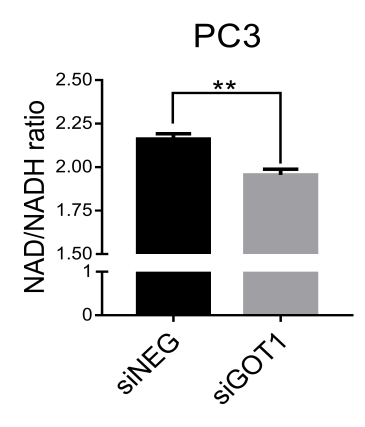

**Supplementary Figure 9.** NAD/NADH ratios in LNCaP and PC3 prostate cancer cell lines. The data from two independent experiments were expressed as mean ± SD.
